# Protective Plant Immune Responses are Elicited by Bacterial Outer Membrane Vesicles

**DOI:** 10.1101/2020.07.24.220160

**Authors:** Hannah M. McMillan, Sophia G. Zebell, Jean B. Ristaino, Xinnian Dong, Meta J. Kuehn

**Author notes:** Corresponding author, Box 3711, Dept. of Biochemistry, Duke University Medical Center, Durham, NC 27710.

## Abstract

Bacterial outer membrane vesicles (OMVs) perform a variety of functions in bacterial survival and virulence. In mammalian systems, OMVs activate immune responses and have been exploited as vaccines. However, little work has focused on the role that OMVs play during interactions with plant hosts. Here we report that OMVs from the pathogenic *Pseudomonas syringae* and the beneficial *Pseudomonas fluorescens* activate plant immune responses that protect against bacterial and oomycete pathogens. OMVs from these two species display different sensitivity to biochemical stressors, which could indicate differences in OMV cargo packaging. Furthermore, our study shows that OMV-induced protective immune responses are T3SS- and protein-independent, while OMV-mediated seedling growth inhibition largely depends on protein cargo. Importantly, OMV-mediated plant responses are distinct from those triggered by PAMP/MAMPs or effector molecules alone. OMVs provide a unique opportunity to study virulence factors in combination and add a new layer of interaction and complexity to host-microbe interactions.

## Introduction

Cells from all kingdoms of life produce extracellular vesicles. Bacteria can use these secreted 40-200 nm biological “packages” to eliminate toxic compounds such as misfolded proteins, facilitate bacterial adaptation to environmental change and stress, and communicate within the bacterial population using insoluble mediators (Orench-Rivera and Kuehn, 2016, Schwechheimer and Kuehn, 2015, McBroom and Kuehn, 2007, Kuehn and Kesty, 2005, Volgers et al., 2018, Florez et al., 2017, Horspool and Schertzer, 2018). In Gram-negative bacteria, these vesicles bud from the outer membrane in a process that does not damage or weaken the bacterial membrane (Kulp and Kuehn, 2010, Schwechheimer et al., 2013, McBroom and Kuehn, 2007, Zhou et al., 1998, Beveridge, 1999). Many factors influence vesicle production, and previous research suggests that production peaks during late log and early stationary phase and increases in response to stress (Kulp et al., 2015, McBroom and Kuehn, 2007, Schwechheimer et al., 2013, Orench-Rivera and Kuehn, 2016, Schwechheimer and Kuehn, 2015, Berleman and Auer, 2013, Pathirana and Kaparakis-Liaskos, 2016). For bacterial pathogens, studies have shown that toxins and virulence factors are preferentially exported in outer membrane vesicles (OMVs), which specifically interact with and are often internalized into host cells (Schwechheimer and Kuehn, 2015, McBroom and Kuehn, 2007, Kuehn and Kesty, 2005, Volgers et al., 2018, Kulkarni and Jagannadham, 2014, Kulkarni et al., 2014, Kulp et al., 2015). OMV-host interaction can benefit the bacteria and contribute to the overall virulence strategy; however, host immune systems also detect OMVs and use them as signals to activate immune responses and overcome infection (Kuehn and Kesty, 2005, Kaparakis-Liaskos and Ferrero, 2015, Ellis and Kuehn, 2010, Acevedo et al., 2014).

Despite the extensive focus on interactions between bacterial OMVs and mammalian hosts, especially in the context of pathogenicity and disease, little is known about the role that OMVs play in the environment and, specifically, their role in plant-pathogen interactions. The sheer abundance of bacteria suggests that OMVs could have major roles in different ecosystems, and already researchers have started to reveal their potential contributions in the marine environment (Biller et al., 2014, Biller et al., 2017, Bahar et al., 2016, Ionescu et al., 2014). Plant-pathogen interactions present a unique opportunity to study OMV’s contributions to bacterial virulence and OMV-mediated interkingdom communication.

Our current understanding of plant-pathogen interactions stems from a large body of work using the model host *Arabidopsis thaliana* and the model bacterial pathogen, *Pseudomonas syringae* pv *tomato* (*Pst*). *Pst* enters the leaf tissue through stomata or wounds in the leaf epithelium and proliferates in the intercellular space known as the apoplast (Katagiri et al., 2002, Xin and He, 2013). This stressful environment, which consists mostly of air and is devoid of nutrients and resources needed for bacterial growth (Xin and He, 2013), can be mimicked *in vitro* using minimal media and has been shown to induce bacterial virulence factor expression (Lam et al., 2014). Plants detect invasion into the apoplast through a combination of extracellular and intracellular defense mechanisms and mount an immune response to clear the pathogen (Spoel and Dong, 2012, Jones and Dangl, 2006, Katagiri et al., 2002, Stael et al., 2015, Xin and He, 2013). Some non-pathogens, including *Pseudomonas fluorescens*, also activate a subset of these plant immune responses at low levels despite their inability to proliferate in foliar tissue (Alfano and Collmer, 2001, Bakker et al., 2007, Pieterse et al., 2014, Weller et al., 2012, Cheng et al., 2017, Haas and Défago, 2005, Iavicoli et al., 2003, Zamioudis and Pieterse, 2011). This low-level plant immune response does not inhibit plant growth significantly, and some of these non-pathogens have even been shown to promote plant growth (Sivasakthi, 2014, Santoyo et al., 2012, Glick, 2012).

A key difference between non-pathogenic commensals and pathogenic bacteria, specifically between *P. fluorescens* and *P. syringae*, is the ability of pathogens to overcome plant defenses using the type III secretion system (T3SS) (Deslandes and Rivas, 2012, Feng and Zhou, 2012, Gassmann and Bhattacharjee, 2012, Guo et al., 2009, Mazurier et al., 2015, Büttner and He, 2009). T3SS studies have been instrumental in identifying and defining the complex pathways in plant innate immune responses, but it is understood that many other bacterial secretion pathways also play a role in plant-microbe interactions. Our growing understanding of virulence in plant pathogenic bacteria has recently expanded to include OMV-mediated secretion and cargo delivery. Proteomic studies have revealed that OMVs from plant bacteria contain plant cell-wall degrading enzymes, components of protein secretion machinery and effectors, nucleic acids known to induce plant immune responses, and a variety of virulence factors (Kulkarni et al., 2015, Sidhu et al., 2008, Solé et al., 2015, Chowdhury and Jagannadham, 2013). Additionally, OMVs from *Xanthomonas campestris* pv. *vesicatoria* and *Xanthomonas oryzae* pv. *oryzae* and virulence factors purified from these OMVs have been shown to trigger immune responses in plants that include callose deposition, increased transcription of pattern recognition receptors, and reactive oxygen species release (Solé et al., 2015, Tayi et al., 2016, Bahar et al., 2016).

Local immune responses to both pathogens and non-pathogens can lead to systemic immune protection (Pieterse et al., 2014). Pathogenic bacteria often induce host expression of the isochorismate synthase 1 (*ICS1*) gene, which encodes an enzyme that catalyzes the production of salicylic acid (SA), a plant immune signal for systemic acquired resistance (Wildermuth et al., 2001, Glazebrook, 2005, Friedrich et al., 1995, Delaney et al., 1995, Cao et al., 1994, Lawton et al., 1996). For non-pathogens, it is widely believed that immune activation is triggered by pathogen/microbe-associated molecular patterns (PAMPs/MAMPs) and that systemic immune induction occurs via SA-independent pathways (Pieterse et al., 2014). In addition to MAMPs, studies have also revealed specific antibiotic, metabolite, and lipoprotein production in *P. fluorescens* strains that elicit local and systemic plant immune responses (Weller et al., 2012, Iavicoli et al., 2003, Tran et al., 2007, Maurhofer et al., 1994). However, little is known about how plants sense and respond to bacterial OMVs, what implications these responses have for the overall immune response to bacteria, and how OMVs play a role in the larger bacterial virulence strategy.

In this study, we show that upon exposure to bacterial OMVs, plants can mount a broad-spectrum immune response against bacterial and oomycete pathogens. This response is conserved for OMVs from a variety of bacterial species, though not all species, and includes a complex range of direct plant immune responses and indirect seedling growth inhibition. Studying these different responses provides unique insight into how plants differentiate between beneficial, commensal, and pathogenic bacterial interactions. Furthermore, our data reveal an exciting new use for OMVs as a tool to uncouple plant growth and defense activation, as well as signaling and immune outcome.

## Results

### *Pseudomonas syringae* and *Pseudomonas fluorescens* outer membrane vesicles

To evaluate OMV production, we isolated OMVs from *Pseudomonas syringae* pv *tomato* DC3000 (*Pst*) and *Pseudomonas fluorescens* Migula ATCC 13525 (*Pf*) grown first in rich and then in minimal liquid media to early stationary phase (McBroom et al., 2006, Chutkan et al., 2013, Lam et al., 2014). We noted that while bacterial cell viability and density were lower in cultures shifted to minimal media compared to those that were mock shifted to complete media, membrane integrity was not compromised (Figures S1A-C), demonstrating that the cells did not die during the shift. We compared OMV production in complete media to production in minimal media by assessing total protein and lipid in the preparations. After controlling for cell growth and culture density, we found that shifting to the minimal media did not alter OMV production in *Pst* or *Pf* (Table S1; Figures S1D-G). Additional characterization by transmission electron microscopy revealed that OMVs produced in complete and minimal media have similar size distributions and morphology (Figures 1A and S1H-K). The size and morphology of *Pst* and *Pf* OMVs are consistent with those reported for OMVs from other bacterial species, with diameters of mainly 50-125 nm (Schwechheimer and Kuehn, 2015, Beveridge, 1999, Kulkarni et al., 2014, Chowdhury and Jagannadham, 2013).

**Figure 1.**
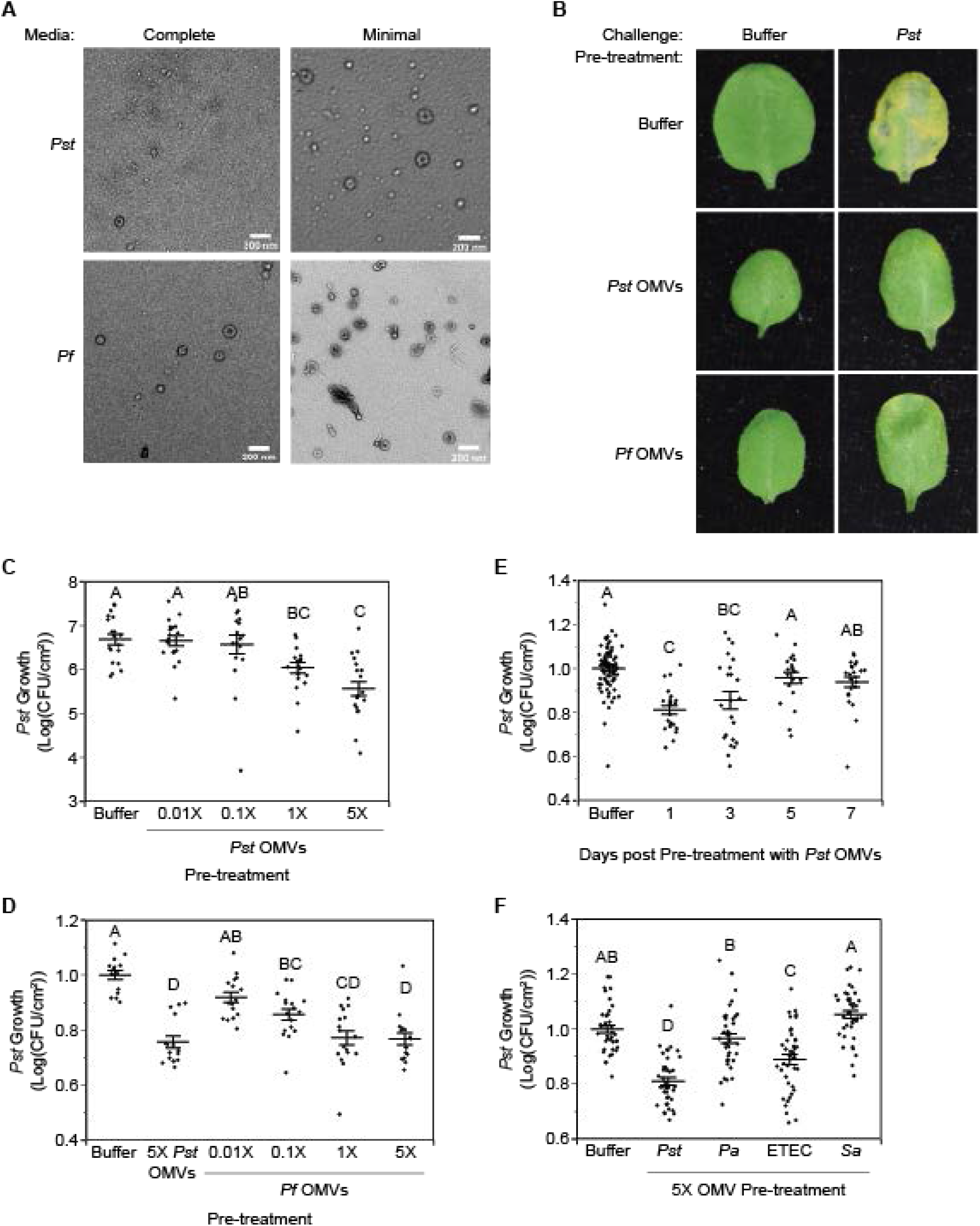
OMV-mediated immune activation protects against bacterial challenge. A. TEM of OMVs isolated from *Pst* (top) or *Pf* (bottom) cultures grown in complete media (left) or shifted to minimal media for 2hr (right). Scale bar: 200 nm. B. Leaves from plants infiltrated with either buffer (left), *Pst* (middle), or *Pf* (right) OMVs from cultures shifted to minimal media for 2hr followed by challenge with either buffer (top) or *Pst* at OD_600_ 0.002 (bottom). C, D. *Pst* growth in plants pre-treated 24hr previously with either buffer or various concentrations of *Pst* (C), *Pf* (D) OMVs from cultures shifted to minimal media for 2hr. Statistics: ANOVA, Tukey HSD. E. *Pst* growth in plants pre-treated with 5X *Pst* OMVs from cultures shifted to minimal media for 2hr. Leaves were pre-treated 1, 3, 5, or 7 days before challenge with *Pst* at OD_600_ 0.002. Statistics: ANOVA, Tukey HSD. F. *Pst* growth in plants pre-treated 24hr previously with either buffer or OMVs/MVs from various species. Mean ± SE. Statistics: ANOVA, Tukey HSD. p<0.05. Conditions not connected by the same letter are statistically significantly different. See also Figure S1 and S2.

### OMV-mediated immune activation protects against bacterial challenge

Given that OMVs elicit immune responses in mammalian systems (Kaparakis-Liaskos and Ferrero, 2015), and that prior exposure to PAMPs enhances the plant immune response to a second challenge (Jung et al., 2009), we hypothesized that exposure to OMVs may have long-lasting immune implications for plants. To test whether OMVs induce protective immune responses, we treated *A. thaliana* leaves with OMVs pelleted from bacterial culture supernatants, waited 24 h, challenged leaves by infecting with *Pst*, then measured bacterial growth after 4 days. Using data collected in Table S1, we calculated the concentration based on protein of OMVs present in a *Pst* suspension at OD600 0.002 and pre-treated with 5X that concentration of OMVs, 13.7 μg/mL. Pre-treatment with *Pst* and *Pf* OMVs resulted in a complete rescue of leaf yellowing in response to *Pst* infection (Figures 1B and S2A). In response to increasing concentrations of OMVs from both *Pst* and *Pf*, we saw a corresponding decrease in bacterial growth (Figures 1C-D). Importantly, OMVs at both the 5X and 1X concentrations significantly reduced *Pst* growth (Figures 1C-D). Further, the leaves remained protected from bacterial challenge at least three days after pre-treatment with *Pst* OMVs, (Figures 1E and S2B).

To determine whether the immune activity was indeed due to OMVs rather than any proteins and small molecules that might co-pellet with OMVs from the culture supernatant, we purified the OMVs further using density gradient fractionation. Previous work from our lab has shown that this method separates OMVs from co-pelleted non-OMV-associated proteins and small molecules (Horstman and Kuehn, 2000, Bauman and Kuehn, 2006). Based on protein and lipid content as well as the evaluation of fraction content by negative-staining electron microscopy, we determined that *Pst* OMVs were present in fractions four through seven of the gradient (Figures 2A and S3A). When each fraction was used to pre-treat plants prior to bacterial challenge, we saw that only the fractions containing OMVs elicited protection against *Pst* (Figure 2B), which suggested that the immune response was in fact OMV-associated. Because no immune activity was found in fractions without OMVs, all subsequent experiments were performed using OMVs pelleted from culture supernatants without density gradient purification.

**Figure 2.**
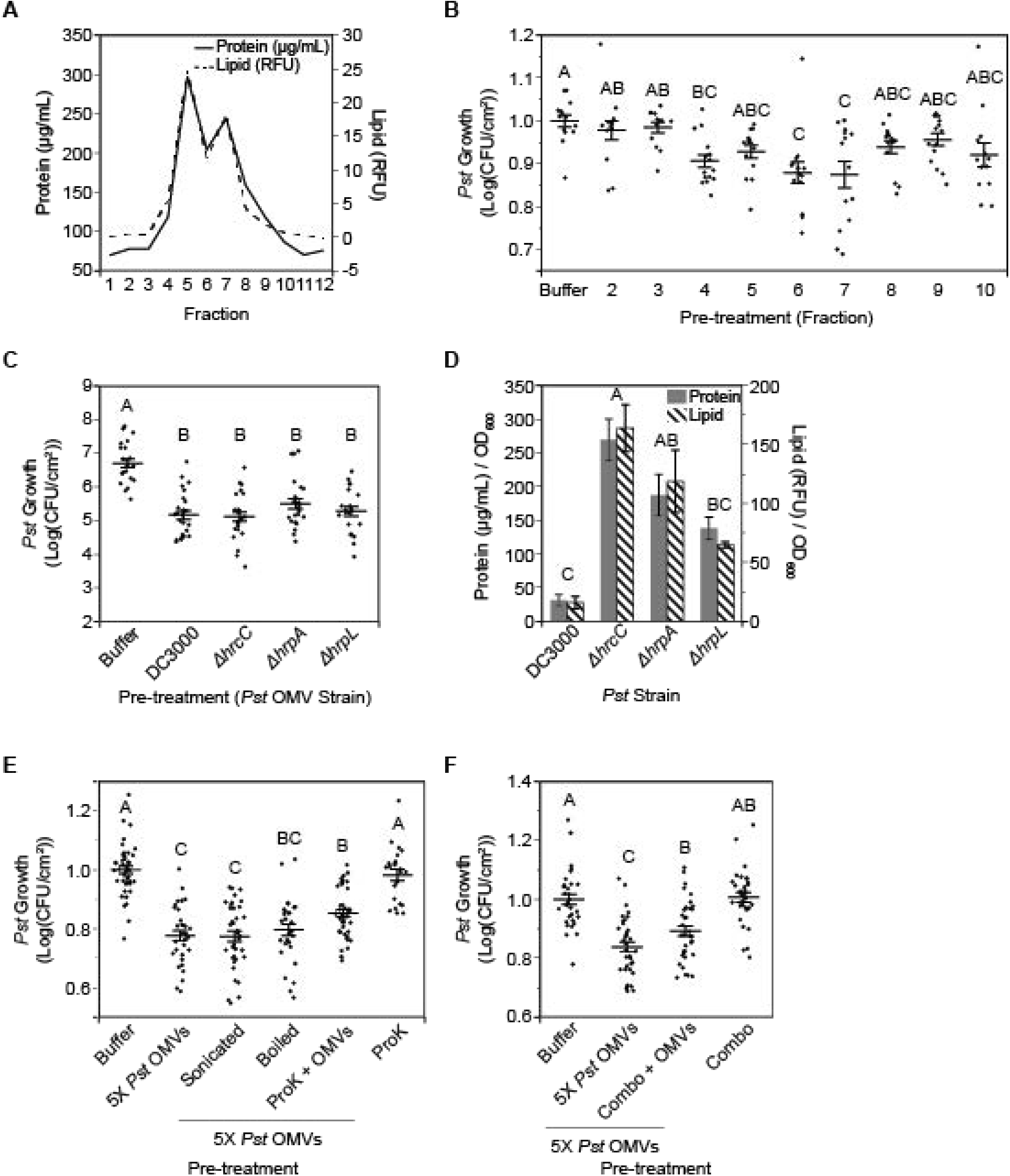
OMV-mediated protection is OMV-associated, Type III Secretion-independent, and withstands biochemical disruption. A. OptiPrep density gradient with *Pst* OMVs from shifted cultures showing distribution of protein and lipid across light (left) to heavy (right) fractions. B. *Pst* growth in plants pre-treated 24hr previously with either buffer or various fractions from the density gradient in (A). Statistics: ANOVA, Tukey HSD. C. *Pst* growth in plants pre-treated 24hr previously with either buffer or 5X *Pst* OMVs from WT, *ΔhrcC*, *ΔhrpA*, or *ΔhrpL*. All OMVs were isolated from cultures grown in complete media and shifted to minimal media for 2hr. Statistics: ANOVA, Tukey HSD. D. OMV production as measured by protein and lipid normalized to culture density. OMVs were isolated from cultures grown in complete media and shifted to minimal media for 2hr. Protein was measured by Bradford assay. Lipid was measured by the lipid dye FM4-64. Statistics: Repeated Measures ANOVA, Tukey HSD. E. *Pst* growth in plants pre-treated 24hr previously with either buffer, 5X *Pst* OMVs, treated 5X *Pst* OMVs (sonicated, boiled, or treated with Proteinase K), or Proteinase K alone. Statistics: ANOVA, Tukey HSD. F. *Pst* growth in plants pre-treated 24hr previously with either buffer, 5X *Pst* OMVs, 5X *Pst* OMVs exposed to a combination (Combo) of treatments (Polymyxin B sulfate [10 μM; 1 h; 37°C], sonication [30 min], Benzonase [20%; 1 h; 37°C], Proteinase K [100 μM; 1 h; 37°C], Tween 20 [2%; 10 min], and boiling [2 h; 100°C]), or the combination treatment alone. Statistics: ANOVA, Tukey HSD. Mean ± SE. p<0.05. Conditions not connected by the same letter are statistically significantly different. See also Figure S3.

Plant immune responses to bacterial infection depend in part on plant recognition of bacterial effector-mediated modifications of plant proteins (Spoel and Dong, 2012, Jones and Dangl, 2006). Both effectors and components of the T3SS machinery have been identified in association with OMVs in various proteomics studies (Kulkarni et al., 2015). To test whether OMV-mediated immune activation was dependent on T3SS factors, we used three well-studied T3SS mutants, *ΔhrcC, ΔhrpA*, and *ΔhrpL*, all of which eliminate the ability of *Pst* to induce leaf collapse and effector-triggered immune responses (Büttner and He, 2009, Roine et al., 1997, Deng et al., 1998, Fouts et al., 2002, Shen and Keen, 1993, Pirhonen et al., 1996, Gopalan et al., 1996, Lindgren et al., 1986). The *ΔhrcC* mutant lacks a core component of the machinery inserted into the bacterial outer membrane, while *ΔhrpA* lacks the pilus protein required to form the needle structure (Deng et al., 1998, Roine et al., 1997). In contrast, *ΔhrpL* lacks a sigma protein required for coordination of various components of the T3SS apparatus and expression of effector proteins; therefore, OMVs from *ΔhrpL* should not contain effectors (Fouts et al., 2002, Shen and Keen, 1993). In response to pre-treatment with OMVs from any of the three T3SS mutant strains, we saw no reduction in protective immune responses as measured by the bacterial challenge assay (Figure 2C). Interestingly, while *in vitro* growth of the three mutant strains did not differ from that of wild type, all three mutants produced significantly more OMVs than wild type (Figures 2D and S3B).

Given that the protective effect of OMVs was T3SS-independent, we next tested whether OMV-mediated protection was at all protein-dependent. Interestingly, treating OMVs with proteinase K or boiling only slightly reduced their protective effect (Figure 2E). In fact, even using a combination of many different biochemical and physical treatments only resulted in slightly reduced protection against bacterial challenge (Figure 2F). For this combined treatment, polymyxin was used to interfere with LPS-mediated immune interactions (Domingues et al., 2012, Cooperstock and Riegle, 1981), sonication was used to disrupt the OMV structure further before treating with benzonase to digest nucleic acid, proteinase K treatment was used to digest any OMV-associated proteins, and boiling for two hours in the presence of the detergent Tween 20 was used to disrupt lipid interactions and denature any remaining proteins. As shown by SDS-PAGE and agarose gel electrophoresis, no detectable proteins or nucleic acids remained in these treated OMV samples (Figures S3C-E). That this combination resulted in only slight reduction in immune activity suggests that protein is not the driving force behind OMV-mediated protection against bacterial infection. It also suggests that the OMV-associated molecules leading to immune activation are highly stable, potentially implicating small molecules.

### *Pst* and *Pf* OMVs protect against oomycete challenge in multiple plant species

To explore how broadly OMV-mediated immune activation was able to protect against pathogens, we tested their effect on oomycete pathogenesis. *Hyaloperonospora arabidopsidis* is a well-studied biotrophic pathogen, requiring a living host to grow and reproduce (Xin and He, 2013, Coates and Beynon, 2010, McDowell, 2014, Kamoun et al., 2015). In *A. thaliana*, pre-treatment with *Pst* OMVs led to a reduction in oomycete growth upon subsequent challenge (Figure 3A).

**Figure 3.**
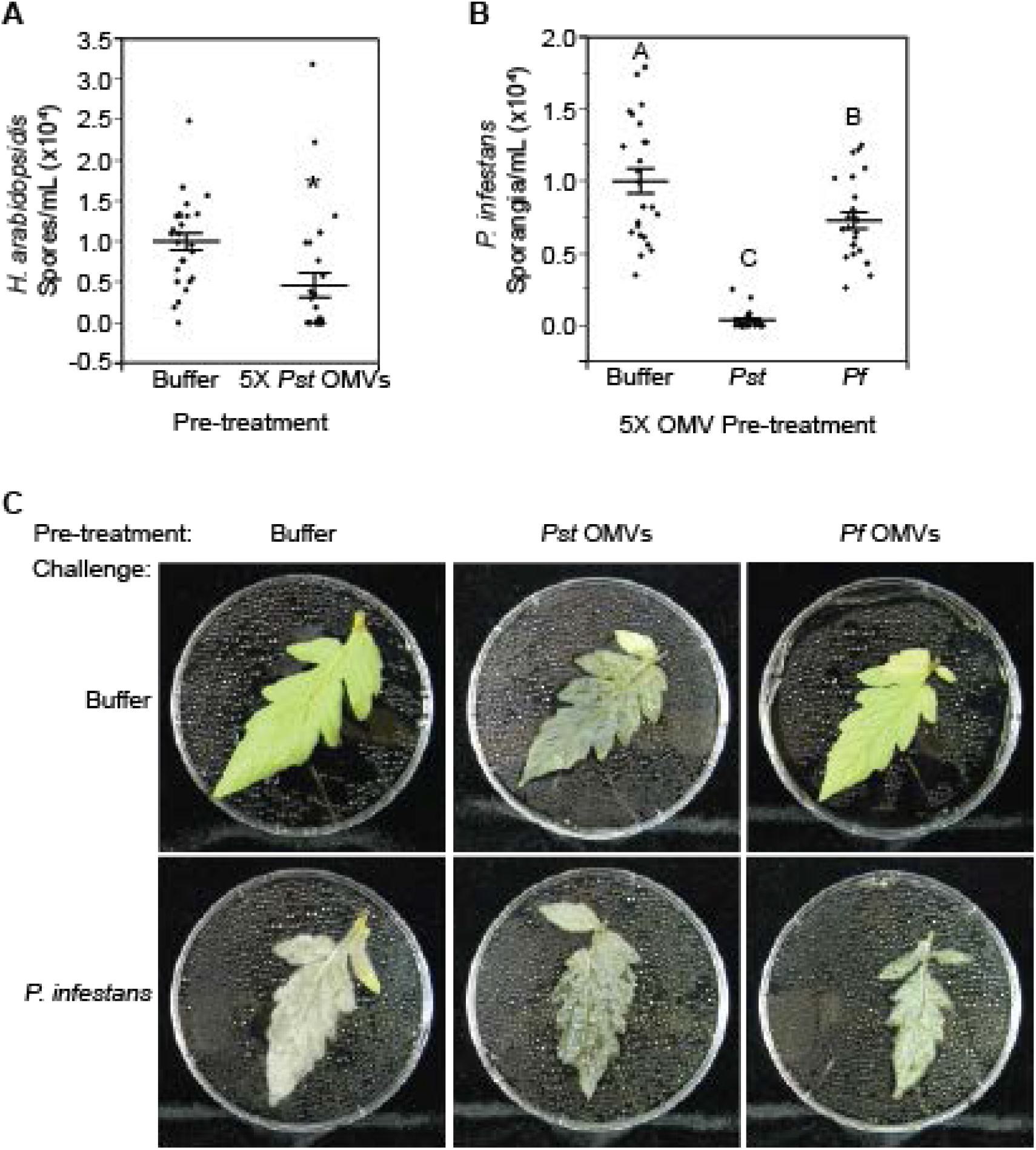
*Pst* and *Pf* OMVs protect against oomycete challenge in multiple plant species. A. *Hyaloperonospora arabidopsidis* spore count in samples isolated from seedlings pre-treated with either buffer or 5X *Pst* OMVs from cultures shifted to minimal media for 2hr. Statistics: Student’s T-test. B. *Phytophthora infestans* sporangia count in samples isolated from detached leaves pre-treated with either buffer, 10X *Pst* OMVs, or 10X *Pf* OMVs from cultures shifted to minimal media for 2hr. Statistics: ANOVA, Tukey HSD. C. Leaves pre-treated with Buffer, *Pst* OMVs, or *Pf* OMVs (top) and challenged with *Phytophthora infestans* (bottom). Mean ± SE. p<0.05. Conditions not connected by the same letter are statistically significantly different. See also Figure S4.

In tomato and potato, another well-studied oomycete pathogen, *Phytophthora infestans* causes devastating disease and crop loss each year (Fry et al., 2015, Kamoun et al., 2015, Ristaino et al., 2019). To test whether bacterial OMVs protect against *P. infestans* infection in its natural host, we pretreated detached tomato leaves of a susceptible cultivar (cv. Mountain Fresh Plus) with *Pst* OMVs, *Pf* OMVs, or a buffer control via vacuum infiltration. Interestingly, pre-treatment with *Pst* and *Pf* OMVs reduced sporangia counts after *P. infestans* challenge (Figures 3B-C and S4A). These data suggest that bacterial OMVs are able to induce protective plant immune responses that improve plant disease resistance during bacterial and oomycete challenge. We note that while *Pst* OMVs did lead to protection against *P. infestans* in tomato, they also led to an interesting water-soaking phenotype that was not observed upon treatment with *Pf* OMVs (Figures 3C and S4A). This same water-soaking was not observed in *A. thaliana* (Figure 1B), leading us to speculate that, as a natural pathogen of tomato, *Pst* may package OMVs with reactive cargo specifically targeting host response pathways in tomato including water-soaking.

### Biochemical and genetic characterization of OMV-mediated growth inhibition

Mounting an immune response requires plants to redirect their resources from growth to defense, resulting in growth inhibition (Albrecht and Argueso, 2017, Lozano-Durán and Zipfel, 2015, Hammoudi et al., 2018, Fan et al., 2014, Huot et al., 2014). Therefore, we hypothesized that OMV treatment would result in growth inhibition as a consequence of immune activation. To test this hypothesis, we treated *A. thaliana* seedlings with increasing doses of *Pst* and *Pf OMVs* and observed *Pst* and *Pf* OMV dose-dependent seedling growth inhibition as measured by seedling weight (Figures 4A-B and S5A-C). Growth was inhibited to a level similar to that of the PAMP flg22 (Figure S5D). Together, these results further support that plants mount an immune response to OMVs.

**Figure 4.**
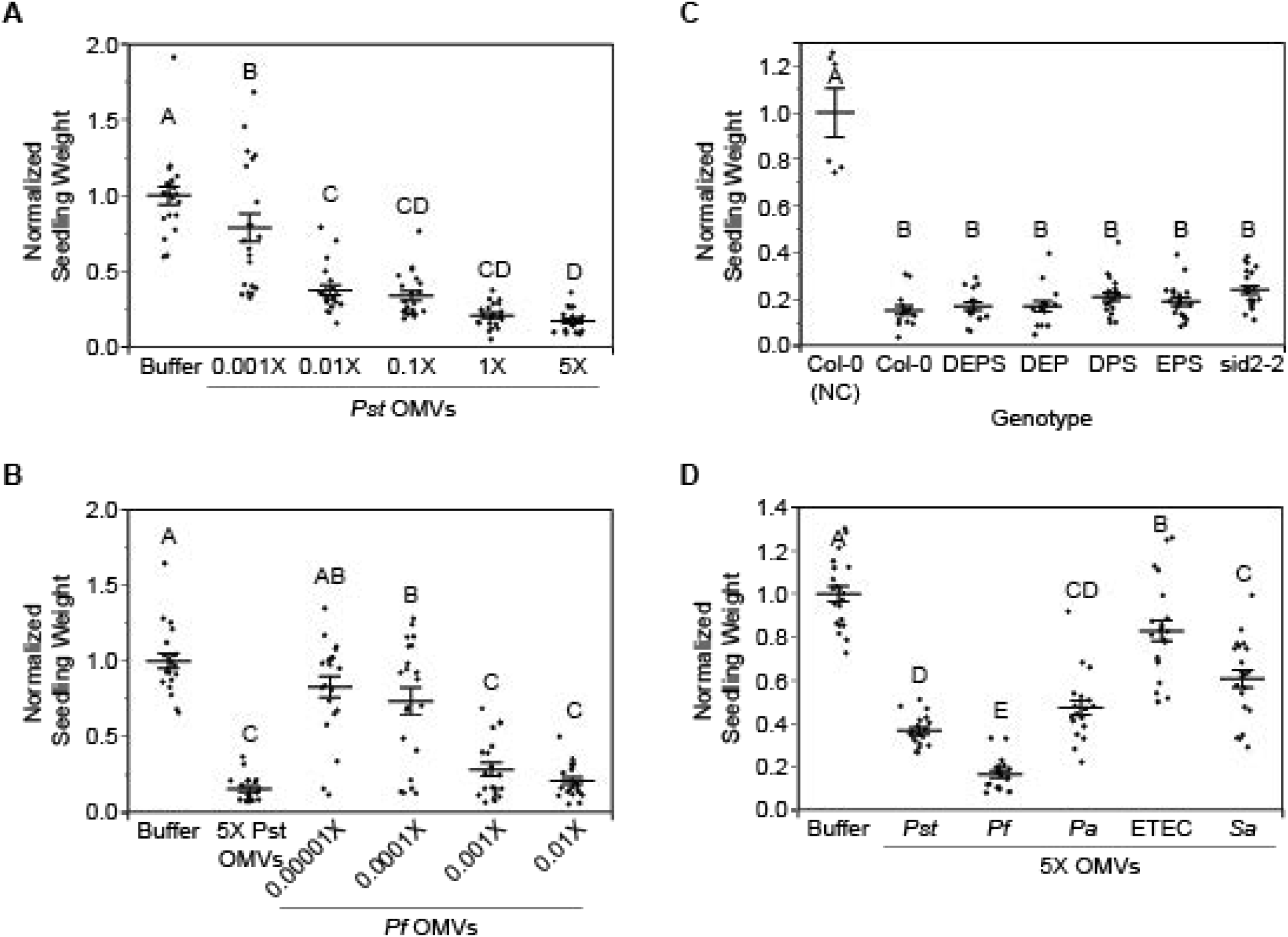
OMV treatment leads to seedling growth inhibition. A, B. Seedling weight 7 days post treatment with either buffer or various concentrations of *Pst* (A) or *Pf* (B) OMVs from cultures shifted to minimal media for 2 h. Statistics: ANOVA, Tukey HSD. C. Hormone mutant seedling weight 7 days post treatment with 5X *Pst* OMVs from cultures shifted to minimal media for 2 h. Weights were normalized to the mean weight of untreated controls from each genotype, then to the mean of Col-0 untreated samples. DEPS: *dde2-2/ein2-1/pad4-1/sid2-2*; DEP: *dde2-2/ein2-1/pad4-1*; DPS: *dde2-2/pad4-1/sid2-2*; EPS: *ein2-1/pad4-1/sid2-2*. Statistics: ANOVA, Tukey HSD. D. Seedling weight 7 days post treatment with either buffer or OMVs/MVs from a variety of bacteria. Statistics: ANOVA, Tukey HSD. Mean ± SE. p<0.05. Conditions not connected by the same letter are statistically significantly different. See also Figure S5.

Using growth inhibition as a convenient and sensitive assay to indicate immune induction we set out to reveal which component(s) of the OMVs were responsible for the protective activities. We applied various biochemical and physical treatments to OMVs and examined their effects on growth inhibition. While disrupting OMV structure via sonication did not alter the growth inhibition phenotype, boiling and proteinase K reduced growth inhibition to the level of the buffer-treated control (Table 1; Figures S5E-F, S5I-K, S5O-P, and S5R). Interestingly, freeze drying and also salt-stripping disrupted the growth phenotype of *Pst* OMVs but not of *Pf* OMVs isolated from minimal media, indicating a difference in packaging despite the similar overall growth inhibition (Table 1; Figures S5E, S5G, S5O, and S5Q). The growth inhibiting activity of *Pst* OMVs from complete media was surprisingly unperturbed by freeze drying but was disrupted by salt-stripping (Table 1; Figure S5J and S5L-M). Similarly, growth inhibition by *Pf* OMVs from complete media remained intact (Table 1; Figure S5J). In notable contrast to the results from the bacterial challenge assays, which showed that protection was protein-independent, these results suggest that seedling growth inhibition is modulated by OMV-associated proteins.

**Table 1.** Summary of growth inhibition and bacterial protection phenotypes in response to treatment or pre-treatment with OMVs.

| Treatment | Growth Inhibition<sup>a</sup>? |  |  |  | Bacterial Protection<sup>a</sup>? |
| --- | --- | --- | --- | --- | --- |
|  | *Pst* ON | *Pst* Shift | *Pf* ON | *Pf* Shift | *Pst* Shift |
| Sonication | Y | Y | Y | Y | Y |
| Salt Strip (NaCl) | N<sup>b</sup> | N<sup>b</sup> | Y<sup>b</sup> | Y<sup>b</sup> | - |
| UV (30 min) | N | N | N | N | - |
| Frozen in LN<sub>2</sub> and Lyophilized | Y | N | Y | Y | - |
| Boiled (100°C, 2 hr) | N | N | N | N | Y |
| Proteinase K ([100µM], 1 hr, 37 °C) | N | N | N | N | Y/N |
| Combined (Polymyxin, Sonicate, Benzonase, Proteinase K, Tween 20, Boil) | N | N | Y/N<sup>c</sup> | N | Y/N<sup>c</sup> |
| Organic Extraction (MeOH and DCM) | Y/N<sup>d</sup> | - | - | - | - |
| Size Exclusion on Extraction Pellet | Y/N<sup>e</sup> | - | - | - | - |

<sup>a</sup>Seedlings (Growth Inhibition) or 3-week-old plants (Bacterial Protection) were treated with OMVs from the indicated strain and condition. OMVs were treated prior to application with the indicated biochemical or physical stressors. Y: activity retained in OMVs; N: activity reduced to level of buffer-treated control; Y/N: activity reduced to a level between OMVs and buffer-treated control. -: activity not tested for these conditions.
<sup>b</sup>Activity retained in stripped supernatant.
<sup>c</sup>Activity was less than 5X *Pst* OMVs and greater than buffer control, but not statistically different from the negative control with treatments alone.
<sup>d</sup>Activity was reduced compared to unfractionated 5X *Pst* OMVs but retained in both fractions.
<sup>e</sup>Activity was reduced compared to unfractionated 5X *Pst* OMV extraction pellet, but only in the fraction containing components larger than 10kDa.

Interestingly, OMV-induced growth inhibition is independent of plant defense hormone signaling. As described previously, abolishing the jasmonic acid, salicylic acid, ethylene, and salicylic acid-independent *pad4* hormone pathways severely compromises plant immune responses (Tsuda et al., 2009). Despite controlling for slight growth differences between the hormone mutants and WT seedlings (Figure S5U), plants lacking these hormone pathways had no effect on the OMVs’ ability to inhibit seedling growth (Figure 4C). This suggests that neither jasmonic acid, salicylic acid, ethylene, nor *pad4* signaling is required for OMV-mediated growth inhibition.

### Bacterial OMVs induce plant *ICS1* expression involved in SA biosynthesis

To assess the plant immune response to OMVs, we utilized *A. thaliana* plants containing a luciferase reporter fused to the *ICS1* gene promoter. Plants infiltrated with *Pst* OMVs showed induced *ICS1* expression, while those treated with *Pf* OMVs did not (Figures 5A-D and S6A). In comparison with bacterial cell-induced *ICS1* expression, we noted that OMVs induced longer lasting but less robust *ICS1* expression (Figures 5A-D and S6A). To confirm that the induced *ICS1* leads to SA accumulation in response to *Pst* OMV treatment, we quantified its production using HPLC. Compared to the buffer treated control, *Pst* OMVs led to increased accumulation of both SAG and SA (Figures 5E-F and S6B). These data suggest that OMVs induce a variety of hormone dependent and independent plant immune responses.

**Figure 5.**
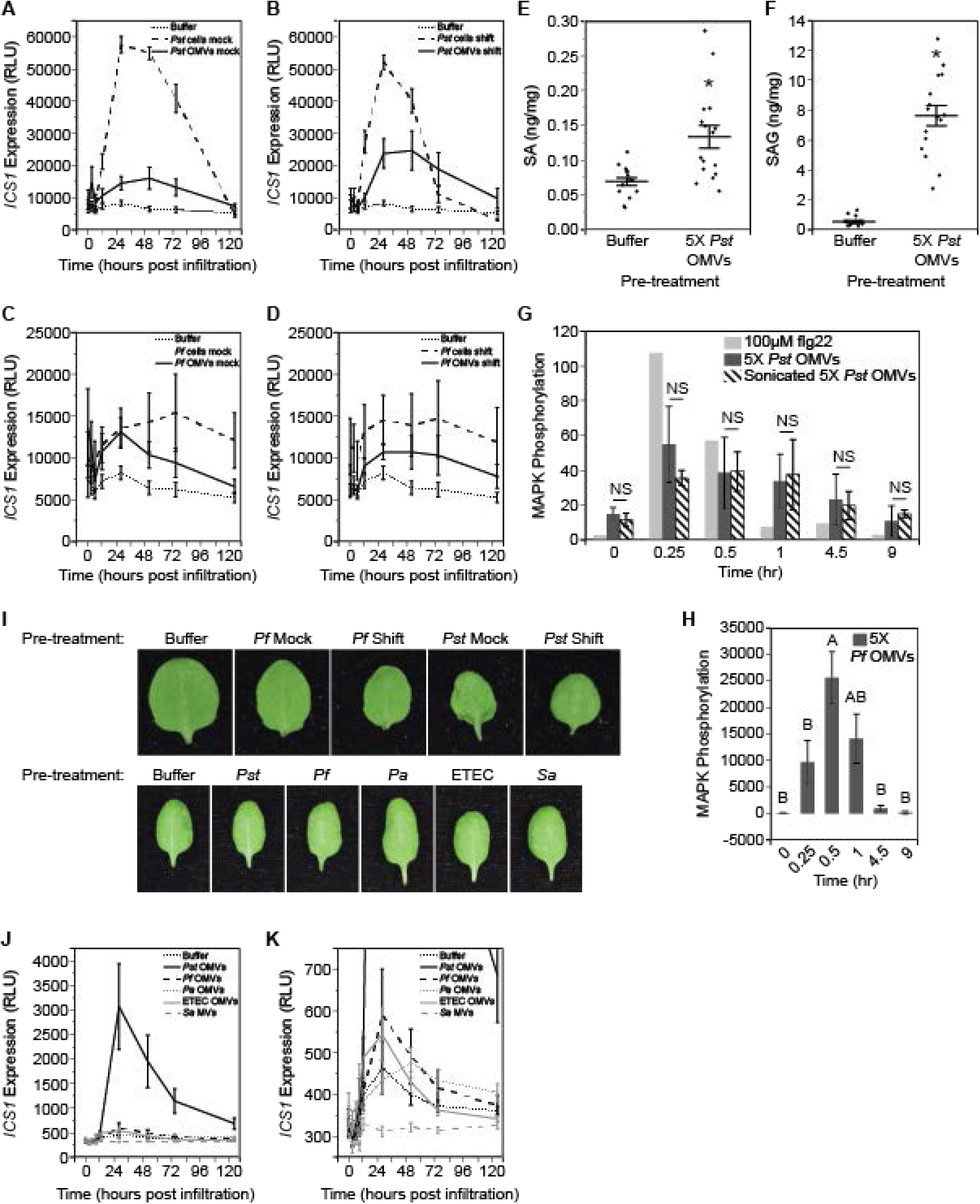
Bacterial OMVs induce plant *ICS1* expression involved in SA biosynthesis. A-D. *ICS1* expression over time from Col-0 *ICS1*:LUC transgenic plants infiltrated with 5X *Pst* OMVs (A, B) or 5X *Pf* OMVs (C, D). OMVs were isolated from cultures grown in complete media and shifted to either (A, C) complete or (B, D) minimal media for 2hr. Statistics: Repeated measures ANOVA, ANOVA subdivided by time point. E, F. HPLC quantification of (E) Salicylic Acid and (F) Salicylic Acid 2-O-β-d-glucose metabolites from leaves infiltrated with *Pst* OMVs from cultures shifted to minimal media for 2hr. Statistics: Student T-test. G, H. MAPK activation over time in response to treatment with flg22, 5X *Pst* OMVs (G), 5X *Pf* OMVs (H), or sonicated 5X *Pst* OMVs (G). Statistics: Repeated measures ANOVA, Student T-test after subdivision by time point. I. Images of infiltrated leaves showing no disease phenotype in *A. thaliana* in association with infiltration of OMVs/MVs from any of the species tested. J. *ICS1* expression over time from Col-0 *ICS1*:LUC transgenic plants infiltrated with OMVs/MVs from various species. Statistics: Repeated measures ANOVA, ANOVA subdivided by time point. K. Zoomed in graph from (J) showing differences in *ICS1* expression among all OMV/MV treatments except *Pst*. Mean ± SE. p<0.05. Conditions not connected by the same letter are statistically significantly different. See also Figure S6.

Similar to the apoplast environment, minimal media induces expression of bacterial virulence factors (Lam et al., 2014). We hypothesized that even though minimal media does not impact OMV production (Table S1; Figures S1D-G), it may influence host reactivity as has been shown in mammalian systems (Orench-Rivera and Kuehn, 2016, Kulkarni et al., 2014, Ellis and Kuehn, 2010). Potentially, OMVs isolated from minimal media cultures might contain more host reactive cargo than those isolated from complete media cultures and would activate stronger immune responses when used to treat plants. However, we found that OMVs isolated from minimal media cultures did not induce significantly stronger or qualitatively different *ICS1* responses than OMVs isolated from complete media, though *ICS1* expression did trend higher with OMVs from minimal media (Figures 5A-D and S6A).

In addition to *ICS1* expression, we found that bacterial OMVs could also activate MAPK phosphorylation known to be involved in PAMP-triggered immunity (Figures 5G-H and S6C) (Tsuda et al., 2013). However, compared to the PAMP signal flg22, OMV-induced MAPK phosphorylation lasted much longer, suggesting that OMVs may activate more complex plant immune responses than a single PAMP signal. The difference in the duration of MAPK signaling has been shown to result in distinct phenotypic immune outputs (Su et al., 2018).

Despite activating plant immune responses, we note that infiltration with OMVs from the tested species or conditions did not lead to water-soaking, leaf yellowing, or collapse in *A. thaliana* (Figures 5I and S6D). Together, these results suggest that *A. thaliana* detects OMVs and mounts immune responses distinct from either the canonical PAMP- or the effector-triggered immune responses, and that *A. thaliana* modulates immune output upon distinguishing OMVs from pathogenic versus beneficial bacteria.

### OMVs from diverse bacteria modulate plant immune responses

To determine whether immune activity and growth inhibition were unique to vesicles derived from model plant bacteria, we isolated OMVs from Gram-negative bacteria *Pseudomonas aeruginosa* (*Pa*), and *enterotoxigenic Escherichia coli* (ETEC), and membrane vesicles (MVs) from the Gram-positive *Staphylococcus aureus* (*Sa*). We then pre-treated leaves with these OMVs/MVs and measured various immune responses or protection against *Pst* challenge. While none of these OMVs/MVs induced *ICS1* expression, OMVs from *Pa*, as well as, to a lesser extent, OMVs from ETEC and MVs from *Sa*, did lead to plant growth inhibition (Table 2; Figures 5J-K and 4D). Interestingly, ETEC OMVs also elicited responses that protected against *Pst* challenge, despite different immune modulation than observed for *Pst* and *Pf* OMVs (Table 2; Figure 1F). These results suggest that protective immune induction is not limited to OMVs from plant-associated bacteria, and that there is cross-protection in response to OMVs from a variety of bacteria.

**Table 2.** Summary of plant immune activation by OMVs/MVs from different bacterial species

| Genus/Species<sup>a</sup> | *Pst* | *Pf* | *Pa* | ETEC | *Sa* |
| --- | --- | --- | --- | --- | --- |
| Visible Phenotype? | N | N | N | N | N<sup>b</sup> |
| Induces *ICS1*? | Y | N | N | N | N |
| Leads to SA Accumulation | Y | - | - | - | - |
| Stunts Seedling Growth? | Y | Y | Y | Y<sup>c</sup> | Y<sup>c</sup> |
| Differential MAPK Phosphorylation? | Y | Y | - | - | - |
| Protects Against Bacterial Challenge? | Y | Y | N | Y | N |
| Protects Against *H. arabidopsidis* Challenge? | Y | Y | - | - | - |
| Protects Against *P. infestans* Challenge? | Y<sup>d</sup> | Y | - | - | - |

<sup>a</sup>OMVs were isolated from the indicated species and applied to plants. Immune activity was measured using a variety of assays. Y: activity present in OMVs; N: activity absent in OMVs; -: activity not tested for these conditions.
<sup>b</sup>Water-soaking phenotype induced in 3-week-old plants at 1-hour post infiltration but returned to normal plant physiology by 3-hours post infiltration.
<sup>c</sup>Seedling growth was stunted to a level between that of buffer and 5X *Pst* OMVs.
<sup>d</sup>Pre-treatment with *Pst* OMVs induces significant water-soaking, possibly destroying leaf tissue and thereby preventing *P. infestans* growth.

## Discussion

Our study shows for the first time that bacterial OMVs elicit plant immune responses that protect against future bacterial and oomycete challenge. Of particular interest, biochemical disruptions of OMVs point to different cargo and likely mixtures of many immunogenic factors, which lead to novel differences in bacterial pathogen-versus beneficial-mediated immune activation. Together, our data provide surprising insight into salicylic acid-independent immune pathways and growth-defense trade-offs, uncover new aspects of OMV-mediated interkingdom communication, and reveal a new layer of complexity in plant-pathogen interactions.

The release by bacteria of OMVs that activate anti-bacterial immune responses may seem counterproductive. However, it should be noted that a parallel situation exists for mammalian bacterial pathogens. Bacterial OMVs are known to elicit strong innate immune responses in mammalian hosts and also harbor highly antigenic epitopes and adjuvanticity that evoke protective and long-term immunity towards the pathogen generating the OMVs (Kuehn and Kesty, 2005, Kaparakis-Liaskos and Ferrero, 2015, Ellis and Kuehn, 2010, Acevedo et al., 2014). A critical artifact of the current reductionist experimental approach is the removal of bacterial cells from the OMV inoculum to study how OMVs behave within plants and interact with the plant immune system. Because of this, these experiments do not address questions about how OMVs and the bacterial cells could work simultaneously, and potentially synergistically, against the plant immune response. Untangling the nuances of how vesicles produced by bacteria contribute to the plant immune response during the various stages of infection is a worthwhile topic for future study and will shed light on a previously unnoticed layer of interaction during natural pathogenesis.

Although both *Pst* and *Pf* OMVs led to protection against bacterial and oomycete challenge (Figures 1C-D, 3A-B, S2A, and S4A), *Pf* OMVs did so independently of *ICS1* expression (Figures 5C-D and S6A; Table 2). The lack of *ICS1* and SA induction is consistent with reports of induced systemic resistance (ISR) by beneficial microbes, which is thought to be mediated by MAMPs (Pieterse et al., 2014, Bakker et al., 2007). For *Pst* OMVs, the *ICS1* induction time course is very similar to that of a PAMP-triggered response (Tsuda et al., 2008, Thilmony et al., 2006, Nomura et al., 2012, Jung et al., 2009). This suggests that the immune activation in response to both *Pst* and *Pf* OMVs was in response to PAMPs/MAMPs or other immune-active molecules contained in or on the OMVs, but also suggests key differences in the exact molecule(s) responsible for protection and in activation pathways. It is important to note that while OMV-mediated immune responses in both *A. thaliana* and tomato resemble those that activate systemic immune responses in plants, systemic protection against *Pst*, *H. arabidopsidis*, or *P. infestans* was not tested in our assays.

Data from the *Pst* T3SS mutants also points towards an immune response like those activated by PAMPs. OMVs from all three mutants protected against bacterial challenge to the same level as OMVs from wild type, which revealed that T3SS machinery and perhaps effectors are not the OMV-associated immune activators (Figure 2C). However, the results from *ΔhrpL* OMVs, which should contain no known effectors, were surprising because effectors have been identified in OMVs from plant pathogens and some of them are potent immune activators (Kulkarni et al., 2015, Kulkarni et al., 2014, Sidhu et al., 2008, Solé et al., 2015, Chowdhury and Jagannadham, 2013, Fouts et al., 2002, Shen and Keen, 1993). Interestingly, all of the T3SS mutants produced significantly more OMVs than wild type, which could be due to fewer connections between the inner and outer membrane, an envelope characteristic found previously to impact OMV production (Schwechheimer et al., 2014, Schwechheimer et al., 2015, Bernadac et al., 1998, Suzuki et al., 1978, Yem and Wu, 1978).

We were particularly surprised by the observation that the protective effect was independent of OMV-associated proteins (Figures 2E-F and S3C-D) while, in contrast, protein cargo was critically important for mediating growth inhibition (Table 1; Figures S5E-F, S5H, S5J-K, S5P, S5R, and S5T). The long incubation with Proteinase K likely led not only to the digestion of all OMV-associated proteins, but also integral membrane proteins, allowing the enzyme to access the OMV lumen and degrade remaining proteins. As a result, these data suggested to us that the cargo of interest in *Pst* OMVs was likely a lipid, small molecule, carbohydrate, or other molecule (Figures 2E-F and S3C-D). Although proteinaceous bacterial effectors are frequently studied for their ability to elicit plant immune responses, many non-protein elicitors have been characterized and are reviewed in several recent publications (Silipo et al., 2009, Morgunov et al., 2017, Erbs and Newman, 2012, Ranf, 2016). Conversely, treating *Pst* or *Pf* OMVs with Proteinase K or boiling OMVs completely eliminated the growth inhibitory effect of OMVs from both strains in both media conditions (Table 1; Figures S5E-F, S5H, S5J-K, S5P, S5R, and S5T). Importantly, this uncouples growth inhibition from immune activation and protection and reveals a novel use for OMVs as probes for plant growth-defense trade-offs upon exposure to complex mixtures of immune active compounds. Consequently, further studies isolating the distinct immune elicitors found in *Pseudomonas* OMVs will likely uncover new aspects of plant immunity.

In comparing *Pst* and *Pf* OMVs, the growth inhibition experiments revealed further differences in plant interaction. Most strikingly, *Pf* OMVs led to much more potent growth inhibition than *Pst* OMVs, as shown by their ability to inhibit seedling growth at a tenfold lower dose (Figures 4A-B and S5A-C). This was surprising since many strains of fluorescent pseudomonads are known to promote plant growth (Bakker et al., 2007, Iavicoli et al., 2003, Sivasakthi, 2014); however, it should also be noted that many of these strains promote plant growth either by helping the plant obtain critical nutrients or by suppressing disease-causing microbes (Berg, 2009, Haas and Défago, 2005, Loper et al., 2007, Glick, 2012).

The activity of OMV preparations from different strains and conditions also varies with application of different biochemical stressors, supporting our hypothesis that these OMVs consist of distinct cargo (Table 1; Figures S5E-T). For example, flash-freezing eliminated the growth inhibition phenotype in *Pst* OMVs from minimal media but had no effect on *Pst* OMVs from complete media (Table 1; Figures S5E and S5L). Intriguingly, flash-freezing also had no effect on *Pf* OMVs from minimal media (Figure S5O; Table 1). This could suggest that in nutrient rich conditions when virulence factor expression is low, *Pst* packages OMVs similarly to those from a beneficial bacterium.

Bacterial OMVs have a wide array of functions in mammalian studies and have been exploited for human benefit through their use in vaccines (Acevedo et al., 2014, Caruana and Walper, 2020). Here, we show that OMVs could have similarly beneficial applications in plant systems and have laid the groundwork for future experimentation by demonstrating that OMVs elicit plant immune responses that protect against bacterial and oomycete challenge. Of particular interest, our results reveal novel differences in bacterial pathogen-versus beneficial-mediated immune activation and OMV packaging, provide a new tool for probing salicylic acid-independent immune pathways and growth-defense trade-offs, and uncover new aspects of OMV-mediated interkingdom communication. By activating many pathways, OMVs elicit a complex immune response that would be difficult for pathogens to adapt to and overcome, lending support for a role for bacterial OMVs in agricultural applications to promote durable resistance. One pressing future direction is to determine the OMV-associated molecule or set of molecules responsible for plant immune activation. While many experiments are needed to determine the precise mechanisms of OMV-mediated plant immune activation, this work provides new perspective on the complexity of plant-bacteria interactions and differences in inter-kingdom communication and immune activation by pathogenic versus beneficial bacteria, which will be critically important in development of new disease management techniques.

## Acknowledgments

We thank the Arabidopsis Biological Research Center for *dde2-2/ein2-1/pad4-1/sid2-2*, *dde2-2/ein2-1/pad4-1*, *dde2-2/pad4-1/sid2-2*, *ein2-1/pad4-1/sid2-2*, and *sid2-2* seeds; P. Zwack for assistance with the *H. arabidopsidis* experiments; A. Saville and J. Hansel for assistance with the *P. infestans* experiments; H. Yoo for help with HPLC experiments; and M.G. Plue and the Duke University Shared Materials Instrument Facility (SMIF) for training and use of the transmission electron microscope.

This work was supported by NSF IOS grant 1931309 and the Duke SOM Core Voucher Program voucher 1272 to M.J.K., the Duke University Medical Center, and grants from the National Institutes of Health (NIH) (NIH 1R35GM118036) and by the Howard Hughes Medical Institute and the Gordon and Betty Moore Foundation (through grant GBMF3032) to X.D. This work was performed in part at the Duke University SMIF, a member of the North Carolina Research Triangle Nanotechnology Network (RTNN), which is supported by the National Science Foundation (grant ECCS-1542015) as part of the National Nanotechnology Coordinated Infrastructure (NNCI). A portion of the work on *P. infestans* was conducted at NC State University with support from the Kenan Institute of Engineering, Science and Technology.

## Author Contributions

H.M.M and M.J.K designed the research. S.G.Z. designed and carried out the MAPK assays and *H. arabidopsidis* challenge experiments and provided substantial feedback on overall research design. J.B.R. supervised and provided feedback on experimentation with *P. infestans*. X.D. supervised *A. thaliana* research, supplied feedback on overall research design, and provided critical feedback on the manuscript. H.M.M. carried out all other experiments and statistical analysis and prepared the figures. H.M.M. and M.J.K. wrote the manuscript with input from all authors.

## Methods

### Bacterial Culture Conditions

*Pseudomonas syringae* pv *tomato* DC3000 was streaked from frozen glycerol stocks onto King’s Broth (KB) agar plates [2% proteose peptone, 8.6 mM K_2_HPO_4_, 1.4% glycerol, 6 mM MgSO_4_, 1.5% agar] supplemented with 25 mg/mL [30.4 mM] Rifampicin and grown for two days at 28°C. Colonies were used to inoculate 50 mL liquid KB media supplemented with Rifampicin and incubated overnight at 28°C with constant shaking. Aliquots of this overnight culture (1 mL) were used to inoculate 0.5-1 L cultures of KB with Rifampicin, which were incubated at 28°C with constant shaking for 17 h to reach stationary phase. Shaking speed was reduced to 150 rpm to reduce foam accumulation. To shift to minimal media, cells were pelleted in a Beckman Avanti J-25 centrifuge (JLA-10.500 rotor; 10,000 x g; 10 min), supernatant was discarded, and cell pellets were resuspended in minimal media and incubated for 2 h at 28°C. Minimal media consisted of minimal salts [20.2 mM KH_2_PO_4_, 4.3 mM K_2_HPO_4_, 3.8 mM (NH_4_)_2_SO_4_, 0.85 mM MgCl_2_, 0.86 mM NaCl, pH to 5 using HCl] supplemented with 0.01% [0.56mM] fructose. *Pseudomonas fluorescens* Migula ATCC 13525 was cultured as above without Rifampicin. Culturing of other bacterial species was as described previously (*P. aeruginosa*, (Esoda and Kuehn, 2019), Enterotoxigenic *E. coli*, (Manning and Kuehn, 2011), *S. aureus*, (Wang et al., 2018).

### OMV Preparation

Vesicles were purified according to published protocols with a few modifications (Bauman and Kuehn, 2006). Cells were pelleted from cultures in a Beckman Avanti J-25 centrifuge (JLA-10.500 rotor; 10,000 x g; 10 min), supernatant was collected and filtered (0.45 μm HV, Millipore Durapore) to remove any contaminating bacterial cells. OMVs were then concentrated using tangential flow (Cole-Parmer MasterFlex) with a 100,000 kDa cutoff filter (Pall Corporation). OMV concentrate was filtered again (0.45 μm HV, Millipore). OMVs were pelleted from the concentrate in a Beckman Avanti J-25 centrifuge (JLA-16.250 rotor; 35,000 x g; 3 h), and resuspended in 1mL minimal media for 1 h at 4°C. OMVs were filtered for sterilization (0.45 μm HV, Millipore Durapore), before pelleting in a Beckman Optima TLX ultracentrifuge (TLA-100.3 rotor; 90,935 x g (max); 1 h). The OMV pellet was resuspended in 100 μL minimal media (overnight; 4°C) before protein and lipid quantitation. OMVs were used for experimentation within the week or stored at −20°C.

For the Bradford assay, concentrated OMVs were diluted 4 times in dH_2_O and 5 μL of this dilution was added to 150 μL Bradford reagent (VWR). Absorbance was measured at 595 nm and compared to a standard curve to calculate protein concentration. Each sample was measured in duplicate. For the FM4-64 assay, concentrated OMVs were diluted 4 times in dH_2_O to a final volume of 20 μL. Diluted OMVs were added to 560 μL of DPBSS buffer [2.7 mM KCl, 1.5 mM KH_2_PO_4_, 200.2 mM NaCl; 4.3 mM Na_2_HPO_4_, 0.5 mM MgCl_2_·6H_2_O, 0.7 mM CaCl_2_] and 20 μL of FM4-64 (100 ng/mL; Invitrogen) and incubated in the dark (37°C; 10 min; shaking). A negative control was prepared by adding 580 μL of DPBSS buffer to 20 μL of FM4-64 (100 ng/mL; Invitrogen) and incubated as the OMV samples. 280 μL of each sample was added in duplicate to a 96-well black clear bottom plate and fluorescence measured in a plate reader (506 nm excitation, 750 nm emission; Molecular Devices SpectraMax).

For density gradient purification of OMVs, 1.5 mL 60% Optiprep (Sigma) in dH_2_O was mixed with 0.5 mL of the pelleted OMV-containing preparation and loaded on top of a 1 mL cushion of 60% Optiprep/dH_2_O at the bottom of a 12.5 mL ultracentrifuge tube. These were followed by layers (2 mL each) of 40%, 35%, 30%, 25%, and 20% Optiprep/dH_2_O. Gradients were centrifuged in a Beckman Optima LE-80K ultracentrifuge (SW 41 Ti rotor; 27,000 x g; 12-14 h; 4°C). Fractions (1 mL) were collected from the top of the gradient. Optiprep/dH_2_O was removed by washing two times in 25mL sterile, dH_2_O in a Beckman Optima LE-80K ultracentrifuge (50.2 Ti rotor; 27,000 x g; 1 hr; 4°C). Pellets were resuspended in 1 mL sterile, dH_2_O, analyzed using Bradford, FM4-64, and TEM, and used in bacterial challenge experiments.

### Sytox Green Assay

Membrane integrity was assessed as previously published (Bonnington and Kuehn, 2016). Five samples were prepared for the Sytox Green assay to measure membrane disruption. Two negative controls were prepared by adding 190 μL of KB or minimal media to 10 μL of Sytox Green (0.1mM, Invitrogen). A positive control was prepared by taking a 200 μL aliquot of cells from a 1L culture grown in KB media, pelleting the cells in a Labnet Prism microcentrifuge (10,000 x g; 10 min), discarding 100 μL of the supernatant, and heating the remaining cell suspension in 10% SDS (100°C, 1 h). To this, 10 μL of Sytox Green (0.1 mM; Invitrogen) was added. Two experimental samples were prepared by taking 190 μL from a bacterial culture grown in KB media or shifted to minimal media and adding 10 μL of Sytox Green (0.1 mM; Invitrogen). In all samples, Sytox Green was added in the dark. All five samples were added to a 96-well black clear bottom plate, incubated (10 min; 37°C; shaking in the dark), and fluorescence measured in a plate reader (500 nm excitation, 550 nm emission; Molecular Devices SpectraMax).

### TEM and Negative Staining

As previously published with some modifications (Manning and Kuehn, 2011), to prepare samples for TEM imaging, metal grids (Electron Microscopy Sciences; 300 mesh copper, formvar/carbon) were cleaned using the Pelco EasiGlow Machine Vacuum and then 10 μL of sample was added. Samples were pipetted onto the grid and let set for 1 min before wicking away excess with a Whatmann filter paper. The samples were stained by applying 10 μL of 2% uranyl acetate, incubated 1.5 min, and excess uranyl acetate removed by wicking with Whatmann filter paper. Grids were stored in petri dishes at room temperature. Samples were imaged using a FEI Tecnai G^2^ Twin Transmission Electron Microscope with a spot size of 2 at a voltage of 120kV.

### *ICS1* Expression

As previously published (Tedman-Jones et al., 2008), seeds from transgenic plants encoding a luciferase reporter fused to the *ICS1* promoter were sown on autoclaved soil with added fertilizer and vernalized for 2 days at 4°C in the dark before transferring to 16 hours of light/8 hours of dark at 28°C. After three weeks, two leaves per plant (leaves 3 and 4) were infiltrated with OMVs between 10 and 10:30 AM and plants were sprayed with luciferin substrate (1 mM). *ICS1* expression was then monitored over time by measuring luminescence in a gel doc (BioRad ChemiDoc XRS+).

### SA/SAG Extraction and Quantification

SA/SAG was extracted and measured essentially as described (Liu et al., 2016, Zheng et al., 2012). To measure metabolite production, about 200 mg of leaf tissue was collected per replicate for each treatment condition and the precise weight recorded. Tissue was collected 52 h post treatment infiltration. Tissue was frozen in liquid nitrogen and ground to a powder with a metal bead in a GenoGrinder (500 strokes/min; SPEX Sample Prep). 600 μL of 90% methanol was added to each sample and vortexed. Samples were then sonicated for 16 min in an ice bath. Next, samples were centrifuged in an Eppendorf 5430 R centrifuge (FA-45-30-11 rotor; 20,800 x g; 5 min) and supernatant was collected and set aside. 600 μL of 100% methanol was added to the samples, vortexed, and sonicated for 16 min in an ice bath for a second time. Samples were centrifuged in an Eppendorf 5430 R centrifuge (FA-45-30-11 rotor; 20,800 x g; 5 min) and supernatant was combined with the supernatant from the initial extraction. Combined supernatant was centrifuged in an Eppendorf 5430 R centrifuge (FA-45-30-11 rotor; 20,800 x g; 5 min) to remove remaining debris. Supernatant from this spin was added to 50 μL of 0.2M NaOH and dried in a Speed-Vac (3 h, 20°C) covered with aluminum foil to avoid prolonged exposure to light. Dried residue was resuspended in 500 μL of 5% trichloracetic acid and sonicated for 16 min in an ice bath. Samples were centrifuged (20,800 x g, 5 min) and supernatant was collected.

Plants produce two forms of SA: the sugar-conjugated form, salicylic acid-2-O-β-D-glucoside (SAG), is produced for storage and the unbound, free SA is thought to be liberated during an infection for use throughout the plant (Dempsey and Klessig, 2017, Summermatter et al., 1995, Blanco et al., 2009). Free SA is considered to be the biologically active form of the hormone (Park et al., 2009, Chaturvedi et al., 2012, Shah and Zeier, 2013, Spoel and Dong, 2012, Dempsey and Klessig, 2012).

Free salicylic acid was extracted 2 times with 500 μL of 1:1 ethyl acetate:cyclopentane, vortexed, sonicated in an ice bath for 16 minutes, and centrifuged at (20,800 x g, 1 min) each time. The upper, aqueous phase was collected from both extractions, combined with 60 μL of HPLC eluent [10% methanol in 0.2% acetate buffer: 0.17% acetic acid, 56.0 mM NaAc, pH 5.5] and dried in a foil-covered Speed-Vac (45 min, 20°C). Samples were dried until 60 μL remained, diluted to 150 μL with HPLC eluent, and analyzed using HPLC with a C18 column (Agilent ZORBAX Eclipse XDB-C18).

Bound salicylic acid (SAG) was extracted by adding 20 μL of 12 N HCl to the combined organic phases of the extraction for free salicylic acid. Samples were heated (80°C; 1 h) and allowed to cool to room temperature slowly. Samples were then centrifuged (20,800 x g; 5 min) and the supernatant was collected. SAG was extracted 2 times with 500 μL 1:1 ethyl acetate:cyclopentane, vortexed, sonicated in an ice bath for 16 min, and centrifuged (20,800 x g, 1 min) each time. The organic phase from both extractions was combined, 60 μL of HPLC eluent was added, and the samples were dried at room temperature in a covered Speed-Vac. Samples were dried until 60 μL remained and then combined with 150 μL of HPLC eluent for HPLC analysis.

Experimental samples were compared to a negative control containing methanol and a standard curve established using dilutions of purified salicylic acid at 0.025, 0.05, 0.1, 0.2, 0.4, 1, 2, 4, and 20 ng/μL. Salicylic acid was quantified by integrating the peaks from a spectrofluorometer (295-305nm excitation/405-407 nm emission).

### Seedling Growth Inhibition

As previously published with some modifications (Xu et al., 2017), seeds were sterilized using isopropyl alcohol and bleach solution prior to plating, and growth inhibition was measured as previously published with some modifications (Navarro et al., 2008). Briefly, seeds were shaken in isopropyl alcohol (70%; 10 min), rinsed with dH_2_O, shaken in bleach (10%; 10 min), rinsed with dH_2_O two times, shaken in dH_2_O (10 min), then vernalized (4°C; shaking in dH_2_O; 2 days). Seedlings were grown vertically on Murashige and Skoog (MS) agar plates [2.3 mM MES, 0.43% MS Basal Salts (Caisson Labs, macro- and micronutrients), pH 5.7 with 1 M KOH, 58.4 mM sucrose, 0.1% MS Vitamins (Caisson Labs), 0.4% agar), at room temperature under 16 h of light/8 h of dark. After 5 days, seedlings were transferred to liquid MS media in 96-well clear plates under sterile conditions. Plates were sealed with micropore tape and seedlings were returned to normal growth conditions. After 2 days, seedlings were treated with 10 mM Mg_2_SO_4_ or dH_2_O buffer control or varying concentrations of OMVs. After 7 days, seedlings were removed from liquid media, blotted on a lint-free tissue to remove excess media, and weighed. For each experiment, 7 plants were grown per treatment. Each experiment was repeated 3 times.

### Bacterial Challenge

As described previously with some modifications (Wang et al., 2006), seeds were sown on autoclaved soil with added fertilizer and vernalized (2 days; 4°C in the dark) before transferring to 16 h of light/8 h of dark at 28°C. After three weeks, two leaves per plant (leaves 3 and 4) were infiltrated with OMVs between 10 and 10:30 AM and returned to the growth room overnight. 24 h later, leaves pre-treated with OMVs were infected with *Pst*. *Pst* was grown for 2 days on KB agar plates at 28°C and suspended in 10 mM Mg_2_SO_4_ to an OD_600_ of 0.002 before infection. Plants were returned to the growth room for 4 days when leaf yellowing started to occur. Using a standard hole punch, discs were taken from each treated leaf and discs from the same plant were ground using a metal bead in 10 mM Mg_2_SO_4_. Samples were then serially diluted and plated in 10 μL strips on KB agar plates. Plates were incubated (2 d, 28°C) and then colony forming units were counted as a measure of bacterial growth.

### OMV Disruption

OMVs were isolated from minimal media as above and then exposed to a variety of disrupting conditions. Separately or in combination, OMVs were treated with polymyxin B sulfate (Sigma-Aldrich) (10 μM, 1 h, 37°C), sonication (30 min, in a water bath), Benzonase (Sigma ≥ 250 units/μL; 20%; 1 h; 37°C), Proteinase K (Sigma-Aldrich, 100 μg/mL,1 h, 37°C), Tween 20 (2%, 10 min; 25°C), or boiling (100°C, 2 h). All OMV preparations were diluted to the 5X concentration (13.7 μg/mL protein) after the disruptions prior to plant infiltration to dilute potentially harmful concentrations of detergent. After diluting for infiltrations, samples contained 0.05% Tween 20. Salt-stripping was performed by treating OMVs with NaCl (2 M, 1 h, 25 °C), pelleting in a Beckman Optima TLX ultracentrifuge (TLA-100.3 rotor; 90,935 x g (max); 1 h), separating the supernatant, and resuspending the stripped OMVs in dH_2_O. For lyophilization, OMVs were flash-frozen in liquid nitrogen and then lyophilized for 1 h. For UV treatment, OMVs were exposed to 254 nm ultraviolet light for 30 min.

### MAPK Activation

As previously published with some modifications (Xu et al., 2017). Phospho-p44/42 MAPK antibody (Cell Signaling #9101) was used at 1:4000 in 5% milk and exposed with SuperSignal West Dura substrate (Thermo Fisher).

### *H. arabidopsidis* Challenge

As described previously with some modifications (Wang et al., 2011), 3 pots per treatment of approximately 50 7-day old Col-0 *A. thaliana* seedlings were dipped into 10 mM MgCl_2_ + 0.05% Silwet L77 or 5X OMVs in 10 mM MgCl_2_ + 0.05% Silwet L77. Gentle vacuum was applied for 3 minutes and released slowly to infiltrate. Plants were covered with a plastic dome and placed in 12 h light/12 h dark growth incubator. 24 h post infiltration, plants were sprayed with 3-5 x 10^4^ fresh spores of *Hyaloperonospora arabidopsidis* NOCO_2_ isolate. Plants were then covered with a mesh dome and returned to growth incubator for 6 days. On the 6th day post infection, plants were covered with a dome and watered heavily to increase humidity, then 10 plants were collected into 1 mL of dH_2_0 for three replicates per pot and vortexed to release spores. Spore density for each replicate was counted 3 times using a hemacytometer. Values are x10^4^ unless otherwise noted.

### *P. infestans* Challenge

Vesicles were purified from *Pseudomonas syringae* pv tomato DC3000 and *Pseudomonas fluorescens* Migula ATCC 13525 and quantified by total protein (Bradford) and relative lipid amount (lipophilic dye FM4-64). Vesicles were diluted in 5 mL of buffer [10 mM MgCl2 + 0.05% Silwet L77] to a concentration of 27.4 μg/mL. Detached leaves of susceptible *Solanum lycopersicum* cv. Mountain Fresh Plus were dipped in either the buffer solution, *P. syringae* vesicles, or *P. fluorescens* vesicles and placed in a petri dish abaxial side up. To infiltrate, a gentle vacuum (25 inHg) was applied for 5 min and released slowly. Leaves were allowed to dry for 6 h at room temperature before transferring under 1.5% water agar plates. Leaves were stored at room temperature under 12 h light/12 h dark for one day prior to late blight inoculation.

*P. infestans* isolate NC 14-1 (clonal lineage US-23) was grown and maintained on detached leaves of *Solanum lycopersicum* cv. Mountain Fresh Plus under 1.5% water agar in ambient laboratory conditions for seven days prior to inoculation. Sporangia were harvested from these leaves by placing the infected leaf in a 15 mL tube with 10 mL dH_2_O and shaking vigorously to release sporangia. Sporangia were quantified using a hemocytometer and sporangial density was adjusted to 1000 sporangia/mL. 500 μL of this solution (500 sporangia) was misted onto the abaxial side of each inoculated leaf using a mist applicator attached to a 15 mL conical centrifuge tube. Conversely, 500 μL of diH_2_O was misted onto the abaxial side of each uninoculated leaf. Leaves were then placed under 1.5% water agar in individual petri plates, wrapped in parafilm, and incubated in plastic bins in ambient laboratory conditions under 12 h light/12 h dark. Sporangia concentration was measured 1-week post-inoculation. Sporangia for each leaf was harvested as above, and the number of sporangia/mL was quantified using a hemocytometer.

### OMV Organic Extraction

Organic extractions were modified from previously published Lipid A preparations (Bonnington and Kuehn, 2016). Using protein to normalize starting amounts, 400 μg/mL of OMVs were diluted to 1 mL in HEPES buffer [50 mM HEPES free acid, pH to 7.4 with 10 mM NaOH] and added to a 12 mL glass vial. 4 mL of 2:1 methanol:dichloromethane was added to the vial, vortexed, and incubated at room temperature for 1 h. Samples were centrifuged in a Sorvall RC 6 Plus centrifuge (SH-3000 rotor; 828.4 x g; 30 min) and the supernatant was collected and set aside. This fraction contained the phospholipids. Pellets were washed with 5 mL of 2:1 methanol:dichloromethane and vortexed to resuspend. Samples were centrifuged in a Sorvall RC 6 Plus centrifuge (SH-3000 rotor; 828.4 x g; 20 min). Supernatant was collected and combined with the supernatant from the first extraction. Pellets were dried with nitrogen and resuspended in 150 μL dH_2_O for use in the seedling growth inhibition assay or 350 μL dH_2_O for the size exclusion fractionation.

### Size Exclusion Fractionation

Pellets from the organic extraction were resuspended in 350 μL dH_2_O. 150 μL of this resuspension was set aside for the seedling growth inhibition assay. The remaining 200 μL was loaded into 0.5 mL Amicon Ultra centrifugal filter devices with a nominal molecular weight limit of either 3,000, 10,000, or 30,000. Samples were centrifuged in a Labnet Prism microcentrifuge (14,000 x g, 30 min), flow-through was collected and diluted to 150 μL in dH_2_O for the seedling growth inhibition assay. Filter tubes were then inverted in the sample collection tubes and centrifuged in a Labnet Prism microcentrifuge (14,000 x g, 15 min) to collect the sample larger than the size cutoff. Flow-through was collected and diluted to 150 μL in dH_2_O for the seedling growth inhibition assay.

### Statistics

All statistical tests were performed using the JMP Pro 14 software using p = 0.05. Means are calculated from at least three experimental replicates. OMV size distributions were calculated using FIJI.

## Supplemental Figures

**Figure S1.**
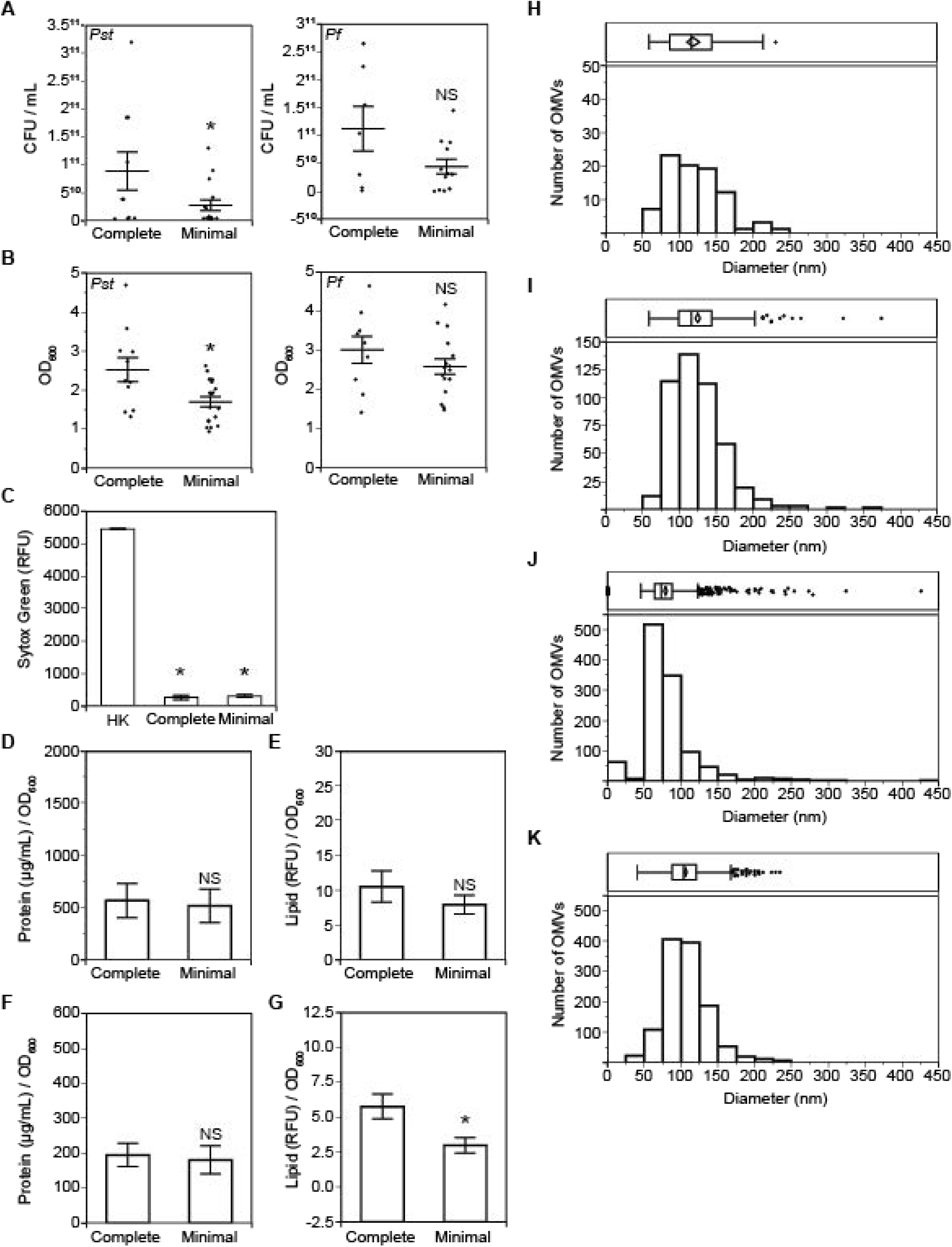
*Pseudomonas syringae* and *Pseudomonas fluorescens* outer membrane vesicles (OMVs). Related to Figure 1. A. CFU of *Pst* (left) and *Pf* (right) in complete versus minimal media. Cultures were grown to early stationary phase in complete media and then shifted to either complete or minimal media for 2hr. Culture aliquots were plated and CFU counted. Mean ± SE. Statistics: Student t-test. B. OD_600_ of *Pst* (left) and *Pf* (right) in complete versus minimal media. Cultures were grown to early stationary phase in complete media and then shifted to either complete or minimal media for 2hr. Mean ± SE. Statistics: Student t-test. C. Membrane integrity of *Pst* in complete versus minimal media as measured by Sytox Green where fluorescence intensity indicates compromised membrane integrity. Cultures were grown to early stationary phase in complete media and then shifted to either complete or minimal media for 2hr. Statistics: ANOVA, Tukey HSD. Mean ± SE. p<0.05. D-G. *Pst* (D, E) or *Pf* (F, G) OMV production as measured by (D, F) protein and (E, G) lipid normalized to culture density. Vesicles were isolated from cultures grown in complete media and shifted to either complete (left) or minimal (right) media for 2hr. Protein was measured by Bradford assay. Lipid was measured by the lipid dye FM4-64. Statistics: Student T-test. Additional tests using repeated measures ANOVA, where CFU and OD_600_, or Protein and Lipid were the repeated measures revealed decreased cell growth in minimal media and no difference in vesicle production, respectively. H-K. Size distribution of OMVs isolated from *Pst* (H, J) or *Pf* (I, K) cultures grown in complete media (H, I) or grown in complete media and shifted to minimal media for 2hr (J, K). The box plot in the top panel summarizes the data where the vertical line is the median, left and right edges of the box are the 1^st^ and 3^rd^ quartile, respectively, left and right whiskers show the 1^st^ quartile minus the interquartile range and the 3^rd^ quartile plus the interquartile range, respectively, confidence diamond shows the mean, where the left and right edges of the diamond are the lower and upper 95% of the mean, respectively, and the points show outliers. Mean ± SE. p<0.05

**Figure S2.**
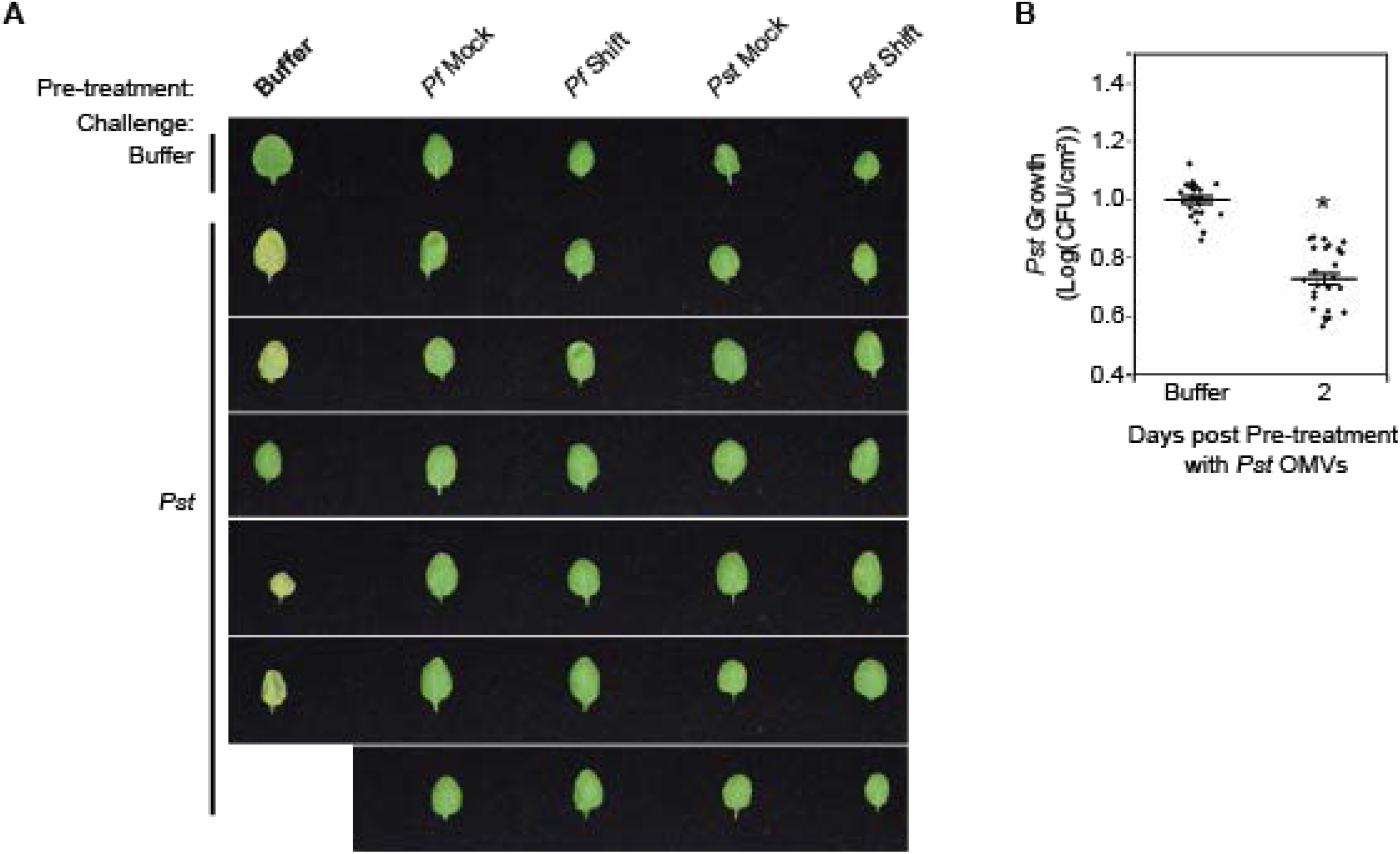
OMV treatment does not lead to *A. thaliana* leaf yellowing and protection lasts at least two days post infiltration. Related to Figure 1. A. Additional images of protection against *Pst* challenge in *A. thaliana* leaves. Leaves were pre-treated with 5X *Pst* or *Pf* OMVs isolated from cultures grown to early stationary phase in complete media and then shifted to either complete (Mock) or minimal (Shift) media for 2hr. Leaves were challenged 24hr post infiltration with *Pst* at OD_600_ 0.002, returned to the growth room for 4 days, and imaged. B. *Pst* growth in plants pre-treated with 5X *Pst* OMVs from cultures shifted to minimal media for 2hr. Leaves were pre-treated 2 days before challenge with *Pst* at OD_600_ 0.002. Statistics: Student T-test. Mean ± SE. p<0.05.

**Figure S3.**
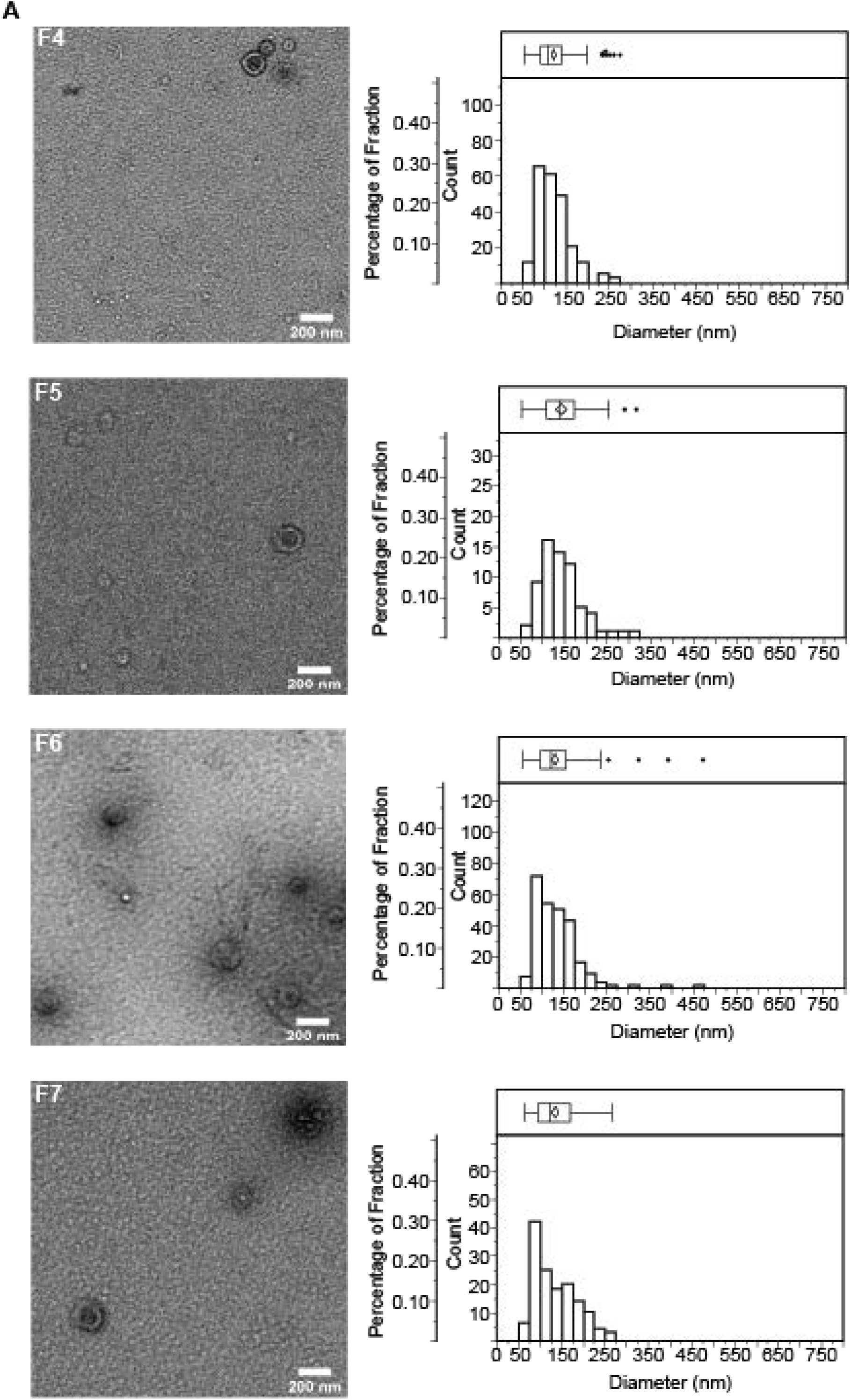
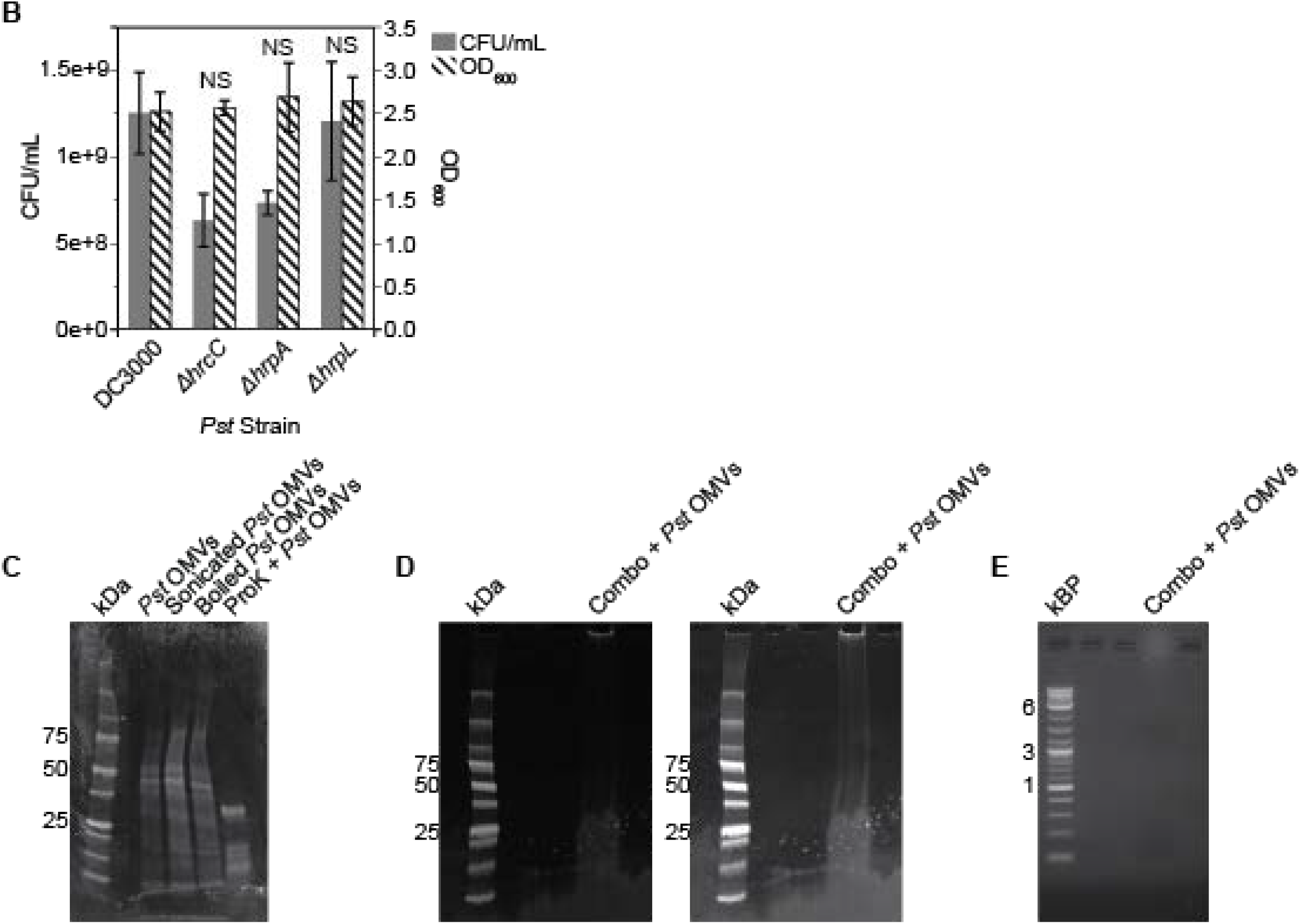
OMVs are present in fractions 4-7 of a density gradient, T3SS mutants display no growth defect *in vitro*, and treated vesicles do not contain detectable protein or nucleic acid. Related to Figure 2. A. TEM images of *Pst* OMVs from OMV-containing fractions of the OptiPrep density gradient with the corresponding size distribution for each fraction. The density gradient was run using OMVs isolated from *Pst* cultures grown in complete media and shifted to minimal media for 2hr. For the size distributions, the box plots in the top panels summarize the data where the vertical line is the median, left and right edges of the box are the 1^st^ and 3^rd^ quartile, respectively, left and right whiskers show the 1^st^ quartile minus the interquartile range and the 3^rd^ quartile plus the interquartile range, respectively, confidence diamond shows the mean, where the left and right edges of the diamond are the lower and upper 95% of the mean, respectively, and the points show outliers. Scale bars: 200 nm. B. CFU / mL (left axis, solid bars) and OD_600_ (right axis, hatched bars) for *Pst* WT and T3SS mutants grown in complete media. Mean ± SE. Statistics: Repeated Measures ANOVA using CFU and OD_600_ as repeated measures for growth. p<0.05. C, D. OMVs from *Pst* cultures grown in complete media and shifted to minimal media for 2hr were treated with sonication, boiling, or Proteinase K (C) or combined (Combo) treatment (Polymyxin B sulfate [10 μM; 1 h; 37°C], sonication [30 min], Benzonase [20%; 1 h; 37°C], Proteinase K [100 μM; 1 h; 37°C], Tween 20 [2%; 10 min], and boiling [2 h; 100°C]) (D) and run on SDS-PAGE to examine protein contents. Proteinase K is 28.9 kDa. The right panel of D shows an overexposed image of the left panel. E. OMVs from *Pst* cultures grown in complete media and shifted to minimal media for 2hr were treated with combined treatment (Polymyxin B sulfate [10 μM; 1 h; 37°C], sonication [30 min], Benzonase [20%; 1 h; 37°C], Proteinase K [100 μM; 1 h; 37°C], Tween 20 [2%; 10 min], and boiling [2 h; 100°C]) and run on a 1% agarose gel.

**Figure S4.**
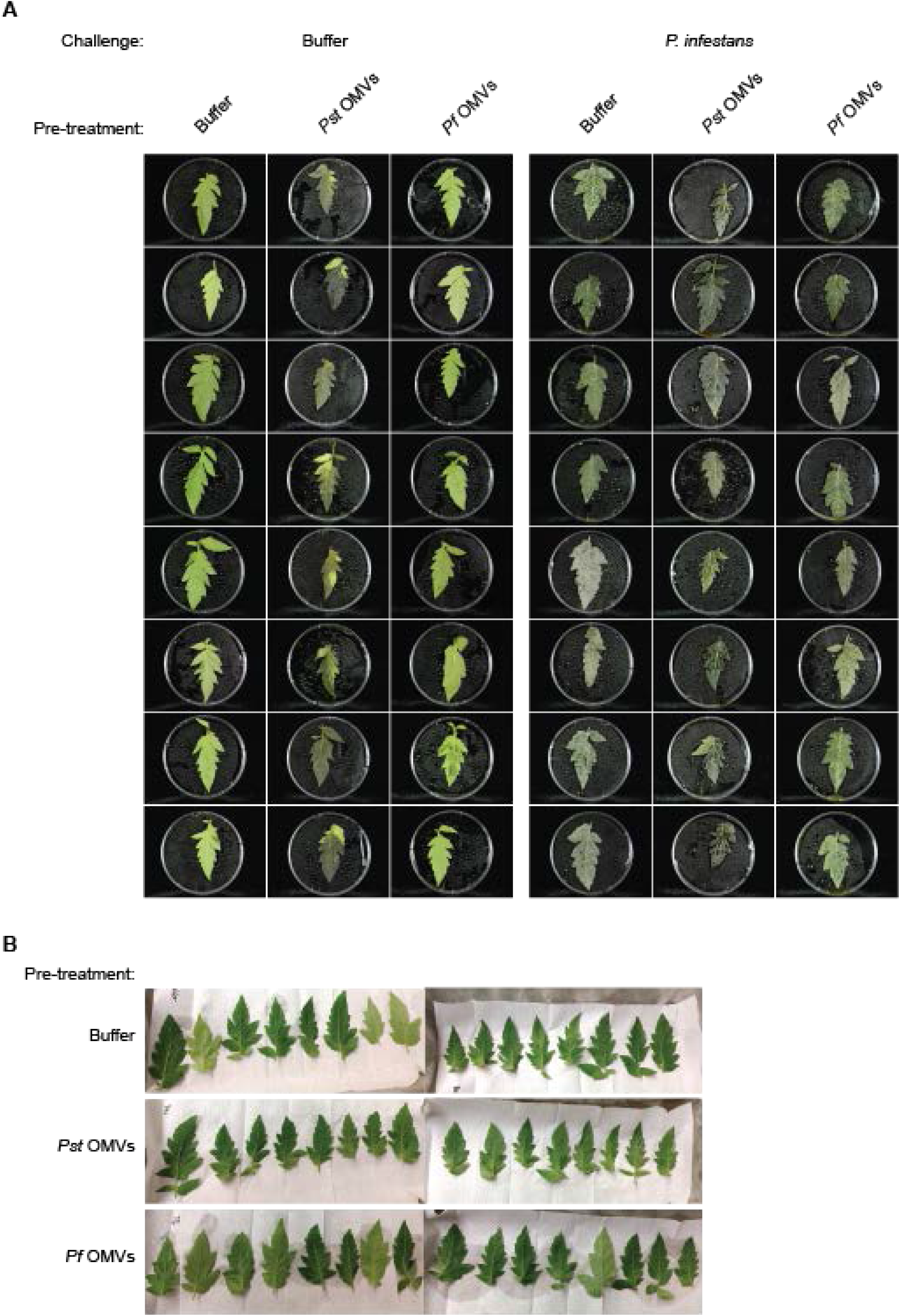
*Pst* OMVs, but not *Pf* OMVs, induce water soaking in tomato, and *Pst* and *Pf* OMVs reduce *P. infestans* growth. Related to Figure 3. A. Additional images of tomato leaves infiltrated with buffer, or 10X *Pst* OMVs or *Pf* OMVs isolated from cultures grown in complete media and shifted to minimal media for 2hr. Leaves were then challenged with buffer (left 3 columns) as a negative control or *Phytophthora infestans* (right 3 columns) and imaged. B. Images of tomato leaves immediately following 10X OMV infiltration from (A).

**Figure S5.**
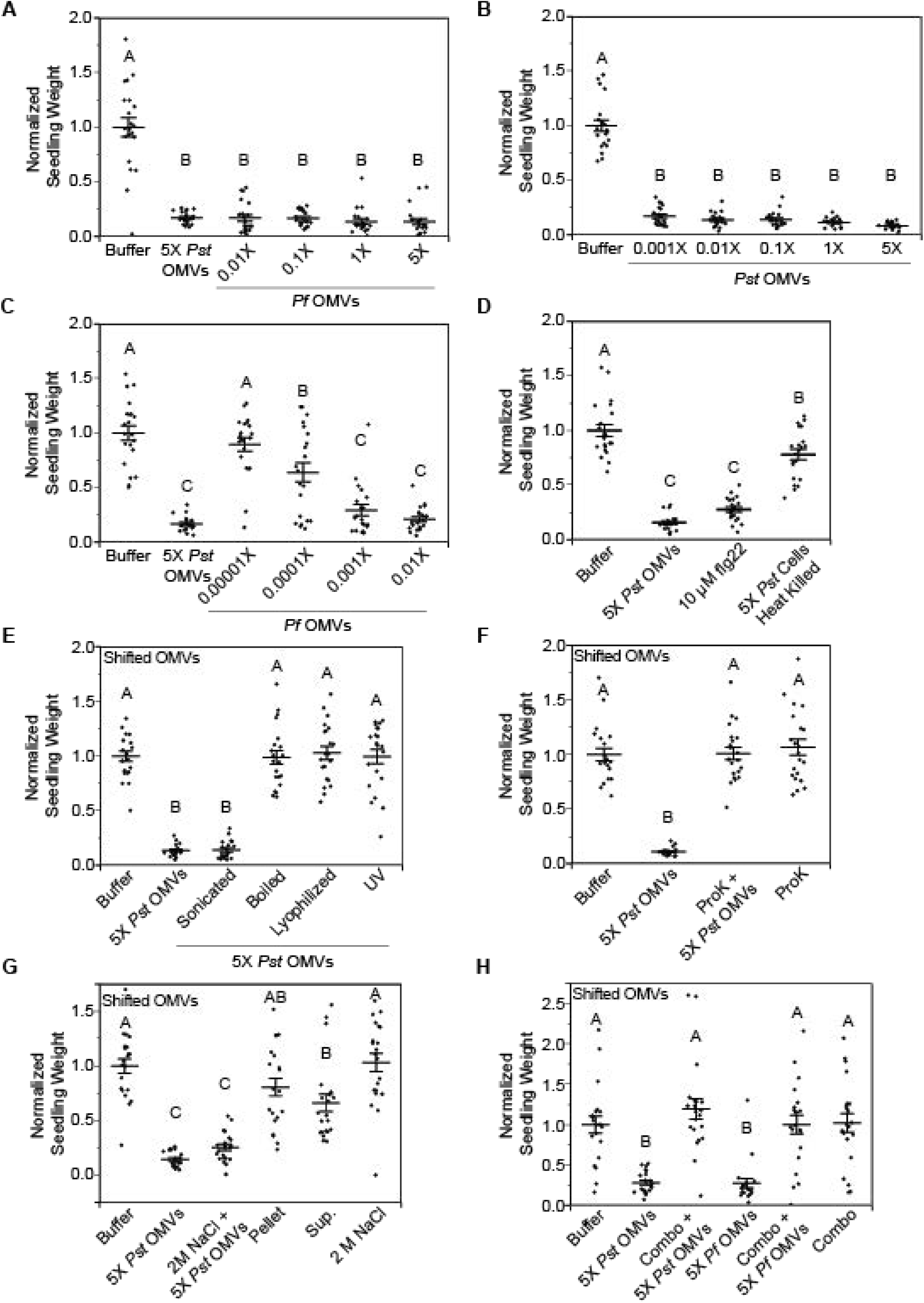
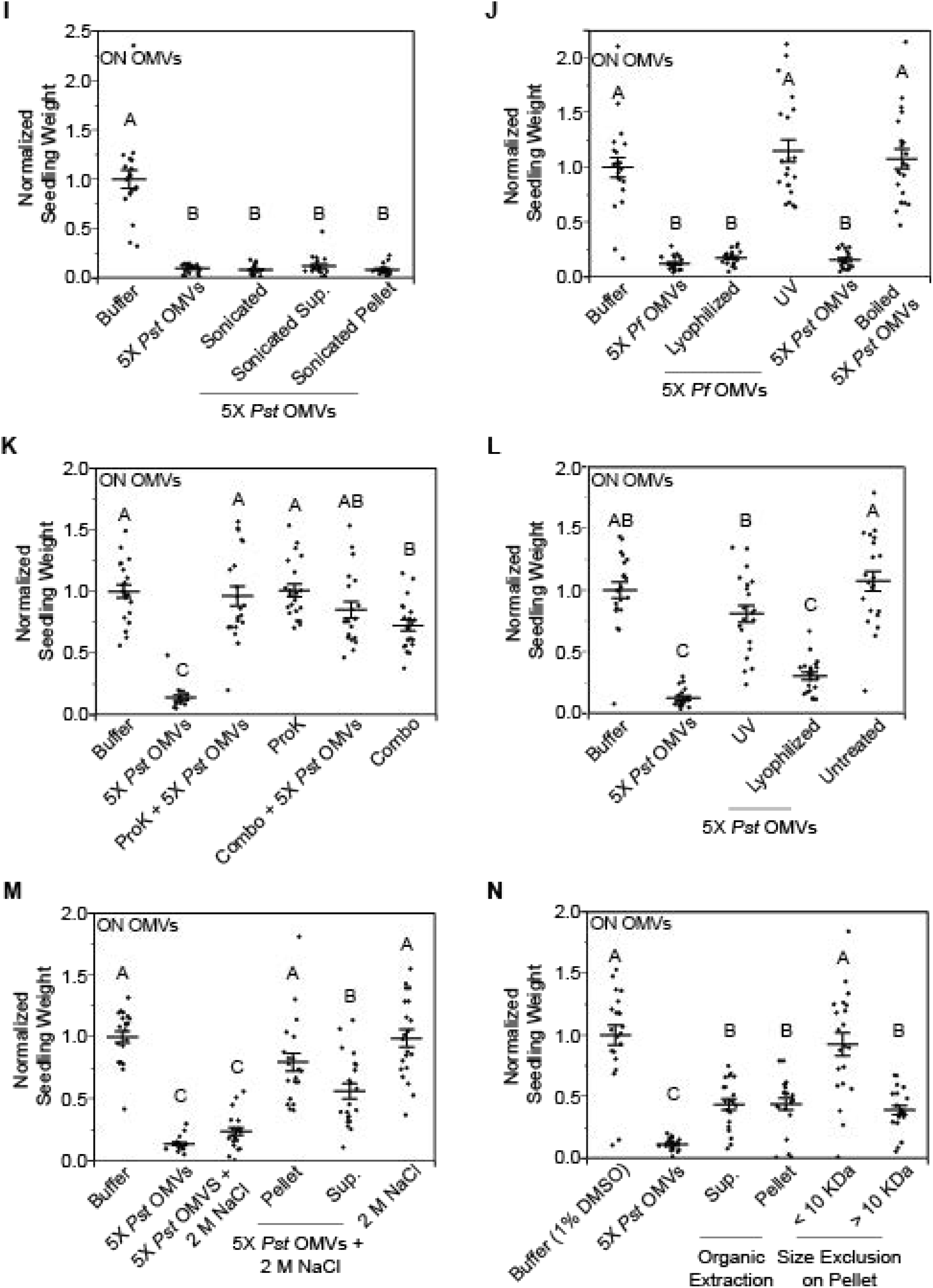
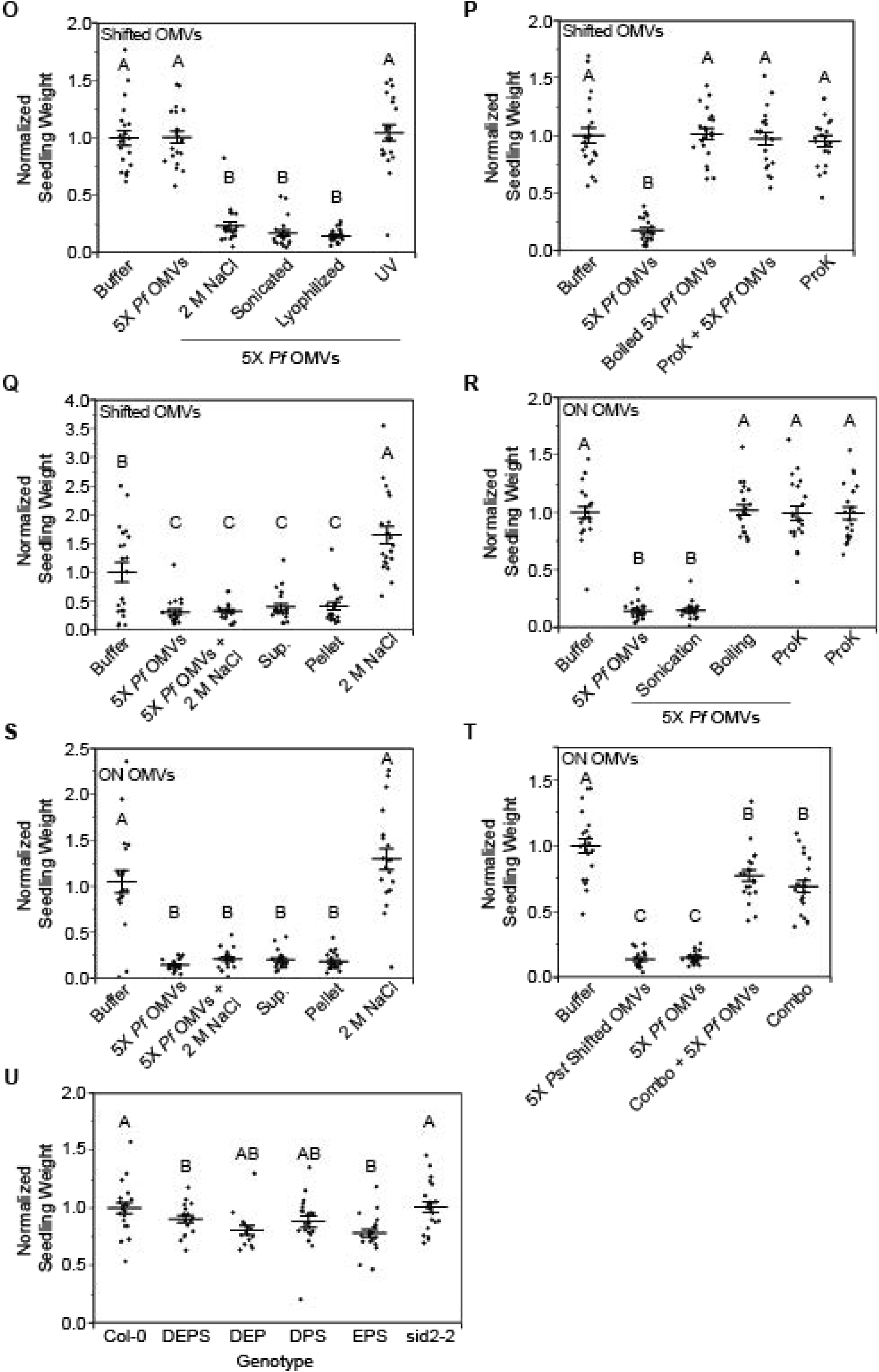
Different biochemical treatments have various effects on the growth inhibition activity of *Pst* and *Pf* OMVs from complete and minimal media. Related to Figure 4. A-C. Seedling weight 7 days post treatment with either buffer or various concentrations of *Pf* OMVs from minimal media (A), *Pst* OMVs from complete media (B), or *Pf* OMVs from complete media (C). Mean ± SE. Statistics: ANOVA, Tukey HSD. D. Seedling weight 7 days post treatment with either buffer, 5X *Pst* OMVs from minimal media, flg22, or heat killed *Pst* cells. Mean ± SE. Statistics: ANOVA, Tukey HSD. E-T. Data in Table 1: Seedling weight 7 days post treatment with OMVs from *Pst* or *Pf* from complete (ON) or minimal (Shifted) media as indicated at the bottom or in the top left of each graph, respectively. Treatments are indicated on the x-axis of each graph. Combined treatment: Polymyxin B sulfate [10 μM; 1 h; 37°C], sonication [30 min], Benzonase [20%; 1 h; 37°C], Proteinase K [100 μM; 1 h; 37°C], Tween 20 [2%; 10 min], and boiling [2 h; 100°C]. (N) Pst OMVs from complete (ON) media were divided into hydrophilic (Pellet) and hydrophobic (Sup.) fractions using a Methanol/Dichloromethane organic extraction. The hydrophilic fraction was further divided into fractions containing particles less than or greater than 10kDa. Mean ± SE. Statistics: ANOVA, Tukey HSD. U. Seedling weight of untreated WT or mutant *A. thaliana* plants 14 days post germination. DEPS: *dde2-2/ein2-1/pad4-1/sid2-2*; DEP: *dde2-2/ein2-1/pad4-1*; DPS: *dde2-2/pad4-1/sid2-2*; EPS: *ein2-1/pad4-1/sid2-2*. Mean ± SE. Statistics: ANOVA, Tukey HSD. p<0.05. Conditions not connected by the same letter are statistically significantly different.

**Figure S6.**
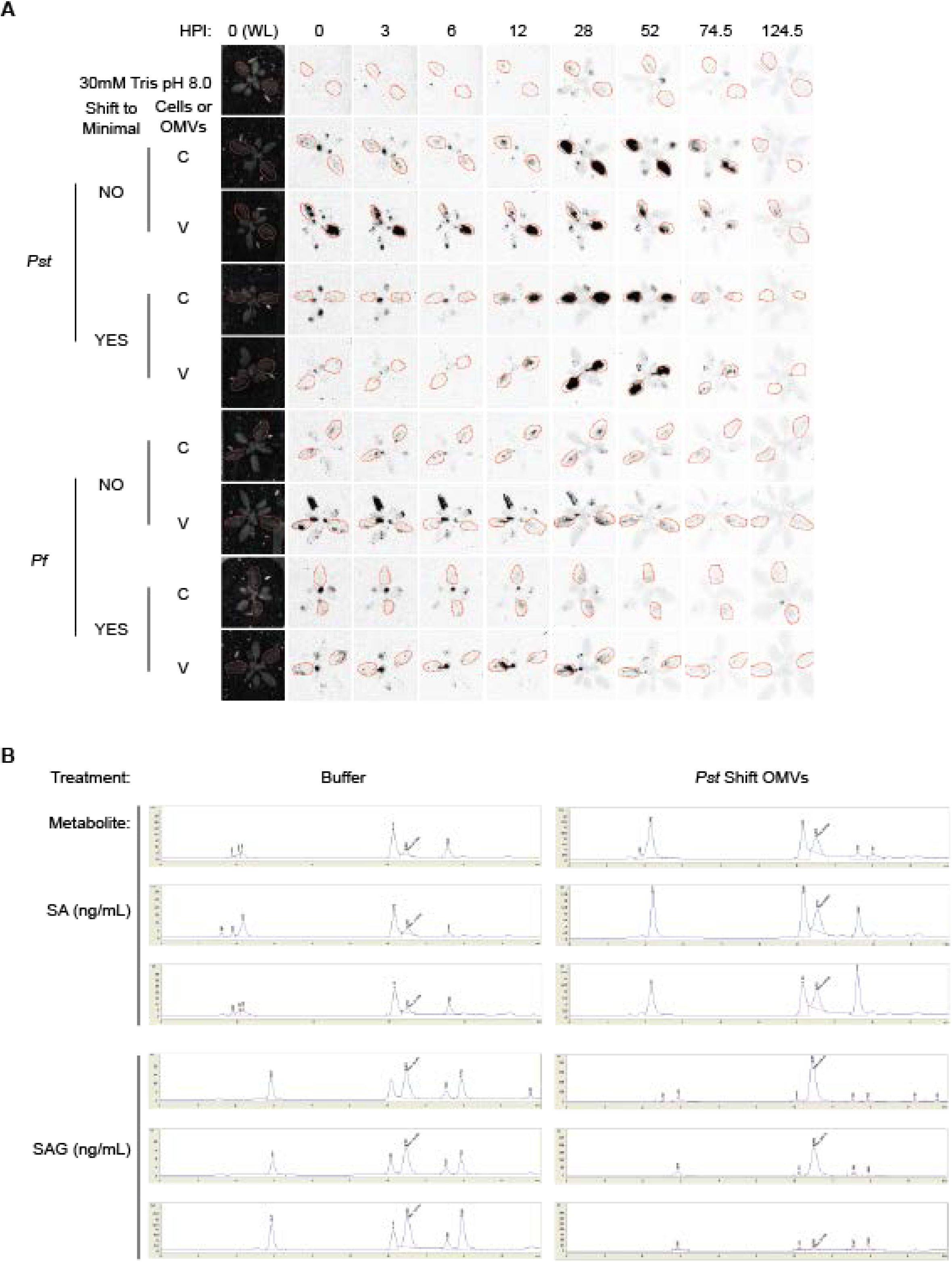
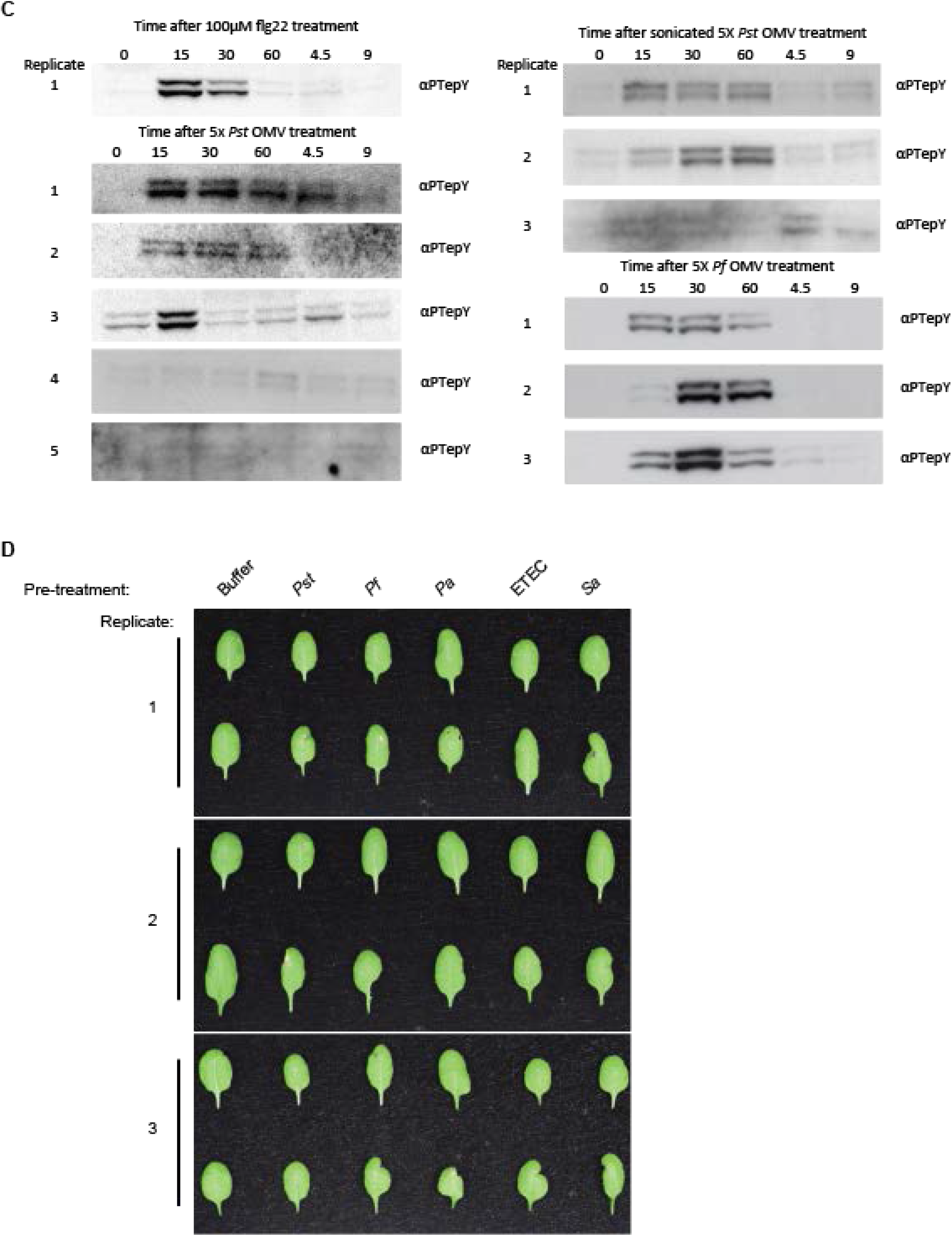
*Pst* OMVs induce ICS1 expression, SA/SAG accumulation, and MAPK activation and no OMVs/MVs tested result in leaf yellowing. Related to Figure 5. A. *ICS1* expression over time as shown by luminescence from Col-0 *ICS1*:LUC transgenic plants infiltrated with OD_600_ 0.002 cells (C) or 5X OMVs (V) from complete (NO) or minimal (YES) media. For shifted cells and OMVs, cultures were grown in complete media and shifted to minimal media for 2hr. Red circles indicate infiltrated leaves. Dark shading indicates luminescence. B. Representative images of HPLC peaks from the SA/SAG purification. Slanted text displays area under the curve for the SA/SAG peak. C. 7-day-old seedlings were treated with flg22, 5X *Pst* OMVs, 5X sonicated *Pst* OMVs, or 5X *Pf* OMVs for the time indicated at the top of each panel. Western blots show phosphorylated MAPK for each replicate. D. Additional images of *A. thaliana* leaves showing no phenotype in response to treatment with OMVs/MVs from various species. Leaves were treated with 5X OMVs/MVs and imaged after 1 week in 16hr light/8hr dark conditions.

## Supplemental Tables

**Table S1.** OMV Yield

| Species | Media<sup>a</sup> | N | Culture Volume (mL) | CFU/mL | OD<sub>600</sub> | Protein (μg/mL) | Lipid (RFU/100μL) |
| --- | --- | --- | --- | --- | --- | --- | --- |
| *Pst* | Complete-Complete | 11 | 500 | 9<sup>10</sup> ± 3<sup>10</sup> | 2.5 ± 0.3 | 1224.8 ± 297.7 | 28.3 ± 9.6 |
| *Pst* | Complete-Minimal | 18 | 500 | 3<sup>10</sup> ± 9<sup>9</sup> | 1.7 ± 0.1 | 773.1 ± 218.9 | 13.3 ± 2.3 |
| *Pf* | Complete-Complete | 9 | 500 | 1<sup>11</sup> ± 4<sup>10</sup> | 3.0 ± 0.3 | 2250.8 ± 436.7 | 67.8 ± 10.0 |
| *Pf* | Complete-Minimal | 16 | 500 | 4<sup>10</sup> ± 1<sup>10</sup> | 2.6 ± 0.2 | 1727.1 ± 349.4 | 33.4 ± 8.6 |

<sup>a</sup>Yield from *Pst* and *Pf* cultures grown in complete media to early stationary phase and shifted to either complete or minimal media for 2 hr.

